# Human Identical Sequences of SARS-CoV-2 Promote Clinical Progression of COVID-19 by Upregulating Hyaluronan via NamiRNA-Enhancer Network

**DOI:** 10.1101/2020.11.04.361576

**Authors:** Wei Li, Shuai Yang, Peng Xu, Dapeng Zhang, Ying Tong, Lu Chen, Ben Jia, Ang Li, Daoping Ru, Baolong Zhang, Mengxing Liu, Cheng Lian, Cancan Chen, Weihui Fu, Songhua Yuan, Xiaoguang Ren, Ying Liang, Zhicong Yang, Wenxuan Li, Shaoxuan Wang, Xiaoyan Zhang, Hongzhou Lu, Jianqing Xu, Hailing Wang, Wenqiang Yu

**Affiliations:** Shanghai Public Health Clinical Center and Department of General Surgery, Huashan Hospital, Cancer Metastasis Institute and Laboratory of RNA Epigenetics, Institutes of Biomedical Sciences, Shanghai Medical College, Fudan University, Shanghai, China; State Key Laboratory of Environmental Chemistry and Ecotoxicology, Research Center for Eco-Environmental Sciences, Chinese Academy of Sciences, Beijing, China; Shanghai Public Health Clinical Center & Institutes of Biomedical Sciences, Shanghai Medical College, Fudan University, Shanghai, China; Shanghai Public Health Clinical Center, Fudan University, Shanghai, China

## Abstract

The COVID-19 pandemic is a widespread and deadly public health crisis. The pathogen SARS-CoV-2 replicates in the lower respiratory tract and causes fatal pneumonia. Although tremendous efforts have been put into investigating the pathogeny of SARS-CoV-2, the underlying mechanism of how SARS-CoV-2 interacts with its host is largely unexplored. Here, by comparing the genomic sequences of SARS-CoV-2 and human, we identified five fully conserved elements in SARS-CoV-2 genome, which were termed as “human identical sequences (HIS)”. HIS are also recognized in both SARS-CoV and MERS-CoV genome. Meanwhile, HIS-SARS-CoV-2 are highly conserved in the primate. Mechanically, HIS-SARS-CoV-2, behaving as virus-derived miRNAs, directly target to the human genomic loci and further interact with host enhancers to activate the expression of adjacent and distant genes, including cytokines gene and angiotensin converting enzyme II (*ACE2*), a well-known cell entry receptor of SARS-CoV-2, and *hyaluronan synthase 2* (*HAS2*), which further increases hyaluronan formation. Noteworthily, hyaluronan level in plasma of COVID-19 patients is tightly correlated with severity and high risk for acute respiratory distress syndrome (ARDS) and may act as a predictor for the progression of COVID-19. HIS antagomirs, which downregulate hyaluronan level effectively, and 4-Methylumbelliferone (MU), an inhibitor of hyaluronan synthesis, are potential drugs to relieve the ARDS related ground-glass pattern in lung for COVID-19 treatment. Our results revealed that unprecedented HIS elements of SARS-CoV-2 contribute to the cytokine storm and ARDS in COVID-19 patients. Thus, blocking HIS-involved activating processes or hyaluronan synthesis directly by 4-MU may be effective strategies to alleviate COVID-19 progression.

## INTRODUCTION

The pandemic of coronavirus disease 2019 (COVID-19) has caused more than 47 million identified cases with more than 1.2 million confirmed deaths worldwide by November 03, 2020 (Dong et al., 2020), and poses an unprecedented global threat. COVID-19 is caused by the severe acute respiratory syndrome coronavirus 2 (SARS-CoV-2) and characterized clinically by fever, dry cough, and shortness of breath (Guan et al., 2020; Mao et al., 2020). Furthermore, COVID-19 may develop acute liver and kidney injury, cardiac injury, bleeding, and coagulation dysfunction (Wiersinga et al., 2020). Severe patients frequently experience acute respiratory distress syndrome (ARDS), causing 70% of deaths in fatal cases (Zhang et al., 2020). Unfortunately, we still lack a specific strategy for COVID-19 treatment.

As a single-stranded positive-sense RNA virus, SARS-CoV-2 is closely related to other highly pathogenic beta-coronaviruses such as severe acute respiratory syndrome coronavirus (SARS-CoV) and Middle East respiratory syndrome coronavirus (MERS-CoV) (Hassan et al., 2020). It has been revealed that SARS-CoV-2 enters into the cell through the binding of spike (S) protein to angiotensin-converting enzyme 2 (ACE2) receptor (Hoffmann et al., 2020; Zhou et al., 2020). SARS-CoV-2 infection activates innate and adaptive immune response (Blanco-Melo et al., 2020), accompanied by elevated inflammation markers (such as C-reactive protein, IL-2R, IL-6, IL-10, and TNF-α) (Chen et al., 2020b). Noteworthily, as the antiviral immunity guardian, T cells are reduced significantly in COVID-19 patients (Diao et al., 2020), which is negatively correlated with survival rates. Besides, increased D-dimer concentration in COVID-19 patients suggests SARS-CoV-2 infection remarkably activates the fibrinolytic system (Levi et al., 2020). However, the molecular mechanism underlying these clinical features caused by SARS-CoV-2 is still elusive.

Accumulating evidence has revealed an interesting phenomenon that both DNA and RNA virus can generate small RNAs different from that of the host cells in the infected cells (Mishra et al., 2019; Shapiro, 2013; Weng et al., 2015), which are called virus-derived small RNAs (vsRNAs) or miRNA-like non-coding RNAs (v-miRNAs). For example, one of the vsRNAs derived from enterovirus 71 (EV71) inhibits viral translation and replication via targeting its internal ribosomal entry site (IRES) (Weng et al., 2014). Conversely, influenza A virus-generated vsRNAs promote virus RNA synthesis (Perez et al., 2010). Except for the regulation of virus replication, vsRNAs also play a key role in regulating host response and disease processes. For instance, repression of LMP2A by miR-BART22 derived from Epstein-Barr virus (EBV) protects the infected cells from host immune surveillance (Lung et al., 2013). Moreover, vsRNAs can mediate the silencing of host genes in *Caenorhabditis elegans* (Guo et al., 2012), and SARS-CoV virus N gene-derived small RNA (vsRNA-N) could enhance the lung inflammatory pathology (Morales et al., 2017). However, little is known about whether SARS-CoV-2 derived vsRNAs participate in the replication of virus and host response in COVID-19 patients.

MicroRNAs (miRNAs) are 19~23 nucleotides (nt) non-coding RNAs (ncRNAs) that primarily regulate post-transcriptional silence via targeting the 3′ untranslated region (3′ UTR) of mRNA transcripts in the cytoplasm (Pasquinelli, 2012). However, our group uncovered that miRNAs located in the nucleus were capable of activating gene expression through targeting enhancer and termed them as “nuclear activating miRNAs (NamiRNAs)” (Xiao et al., 2017). Consistent with our findings, Phillip A. Sharp et al. deciphered the interaction between super-enhancers (SEs) and miRNA networks (Suzuki et al., 2017). Significantly, SARS-CoV-2 was predicted to be enriched in the nucleolus by a computational model named RNA-GPS (Wu et al., 2020). Besides, numerous unknown transcripts have been identified from the architecture of SARS-CoV-2 transcriptome in infected Vero cells (Kim et al., 2020), which may serve as the precursor miRNAs (pre-miRNAs). Therefore, these findings imply that SARS-CoV-2 may generate vsRNAs that function in host cells.

Here, we identified five conserved fragments with high similarity across different primates and termed them as “human identical sequences (HIS)” by comparing the genomic sequences of SARS-CoV-2 and humans. Further bioinformatics analysis indicated HIS embedded in SARS-CoV-2 could potentially be virus-derived small RNAs. HIS-SARS-CoV-2 RNA could directly bind to the conserved human DNA loci *in vitro.* Besides, these virus fragments containing HIS can increase the H3K27 acetylation (H3K27ac) enrichment at their corresponding regions of the human genome in different mammalian cells and activate the expression of adjacent and distant genes associated with inflammation. Notably, HIS can also activate *hyaluronic acid synthase 2* (*HAS2*) and increase the production of hyaluronic acid, which were further verified in COVID-19 patients’ plasma and proven to be correlated with the severity and clinical manifestations of SARS-CoV-2 infection. Hyaluronan inhibitor treatment downregulated the hyaluronan level and thus emerged as a potential therapeutic strategy for COVID-19 patients.

## RESULTS

### Identification of Human Identical Sequences (HIS) in SARS-CoV-2 Genome

Conserved and regulatory elements have been revealed in the viral genome (Bernard, 2013; Morales et al., 2017). Therefore, we hypothesized that nucleic acid sequence of SARS-CoV-2 might interact with host genome as pathogenic factors. To investigate the underlying interaction between SARS-CoV-2 and the host, we analyzed the sequence conservation between the genome of SARS-CoV-2 (accession number NC_045512) and human (GRCh38/hg38). Considering that the shortest regulatory RNAs derived from human genome are 19~21 nt miRNAs; thus, we set the length range of conserved sequences as greater than 20bp and the matching rate as 100%. Surprisingly, we identified five fully conserved sequences in the SARS-CoV-2 genome (Figure1A), which were collectively termed as "Human Identical Sequences (HIS)". HIS-SARS-CoV-2-1 (abbreviated as “HIS-SARS2-1”) and HIS-SARS2-2 were located in chr3 (Figure1B), while HIS-SARS2-3, HIS-SARS2-4, and HIS-SARS2-5 in chr5, chr18, chrX, respectively (FigureS1A).

**Figure 1.**
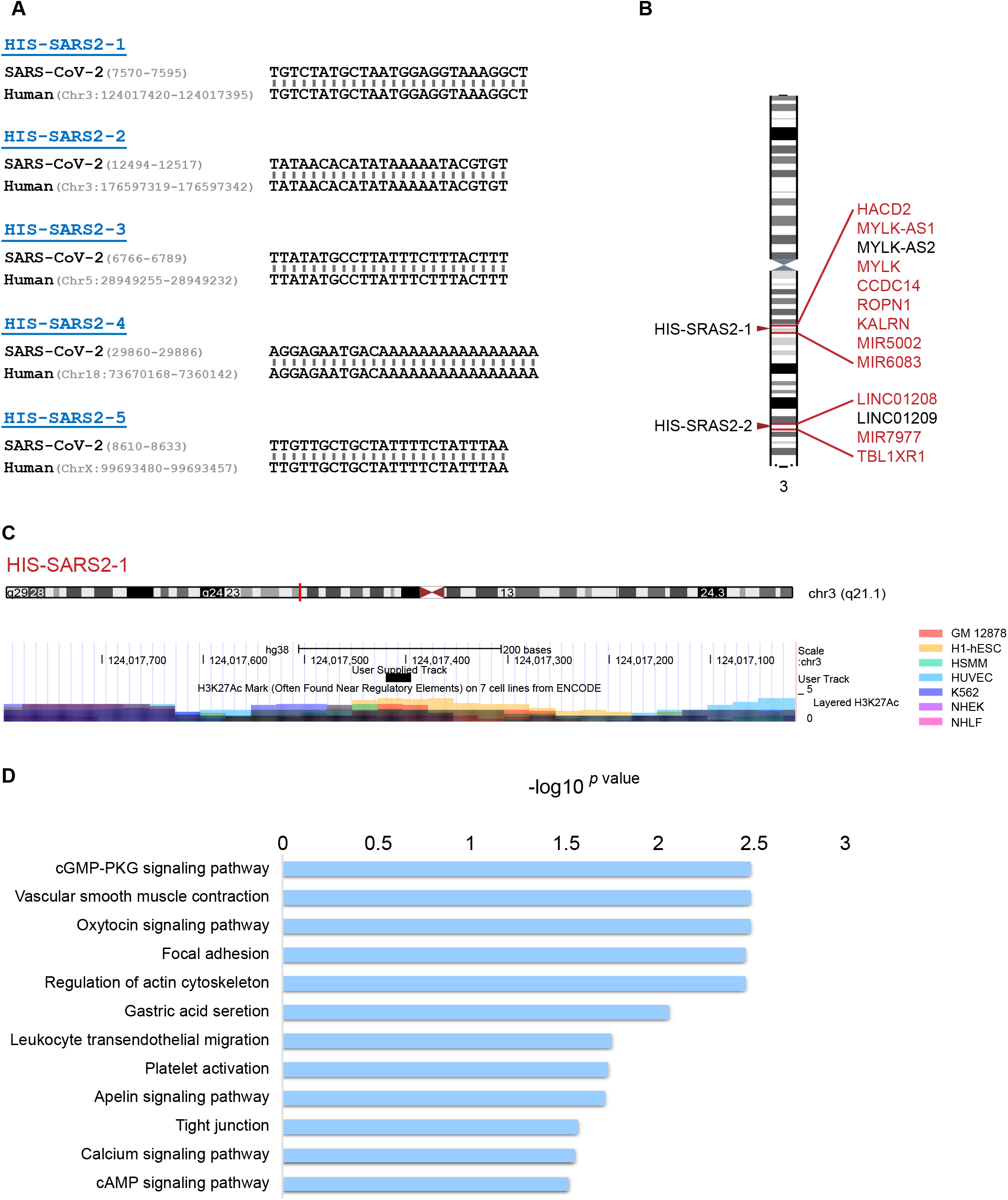
Identification of Human Identical Sequences (HIS) in SARS-CoV-2. **(A)** The sequences of five HIS identified in SARS-CoV-2. In each panel, the upper sequence indicates SARS-CoV-2 genome, while the lower indicates human genome. Numbers in parentheses indicate the location of HIS in the corresponding genomes. HIS, human identical sequences. Chr., chromatin. **(B)** The location of the identical sequence of HIS-SARS2-1 and HIS-SARS2-2 in human chromatin 3 and their surrounding genes. All genes within ± 500 kb of these loci are listed. Among them, inflammation- or immunity-related genes, which are defined by text mining (search combining keywords “gene name + inflammation” or “gene name + immunity”) in PubMed and Google Scholar, are written in red. The searched results were further confirmed by literary research. **(C)** The distribution of enhancer marker H3K27 acetylation (H3K27ac) across the identical sequence of HIS-SARS2-1 in human genome in seven human cell lines. The upper panel illustrates the corresponding chromatin and the lower panel illustrates H3K27ac enrichment, within which the location of the identical sequence of HIS-SARS2-1 is marked in red line and black block, respectively. **(D)** Kyoto Encyclopedia of Genes and Genomes (KEGG) pathway enrichment analysis of genes within ±500kb of the identical sequences of HIS-SARS2 in human genomes. The *x*-axis indicates the degree of KEGG pathway enrichment by rich factor. The *y*-axis indicates the functional terms.

To acquire a better understanding of the potential function of HIS, we interrogated the characteristics of the identical targeted sequences of HIS-SARS2 in human genome. Clearly, we found that all these human genomic loci were widely featured with H3K27ac, the well-known marker of enhancer (Figure1C & FigureS1B), suggesting that HIS-SARS2 may function as a regulator with host enhancer. As enhancer elements embedded in the genome generally regulate their neighboring as well as distant genes, we extended the genomic scope and found there were plenty of well-recognized genes near the targets of HIS-SARS2, including cytokines genes (Figure1B), which might explain why most severe COVID-19 patients are characterized by cytokine storm, the main cause for ARDS that leads to death(Huang et al., 2020). Furthermore, the Kyoto Encyclopedia of Genes and Genomes (KEGG) pathway analysis of the neighboring (±500kb) genes of HIS showed that they were enriched in cGMP-PKG signaling pathway and muscle contraction (Figure1D), consistent with the proposals that modulation of cGMP-PKG by the inhibitor of its upstream regulator phosphodiesterase 5 (PDE5), which is highly expressed in airways and vascular smooth muscle, could be a potential treatment for COVID-19 patients (Giorgi et al., 2020). Gene Ontology (GO) analysis showed that they were enriched in vascular smooth muscle contraction and protein import into the mitochondrial matrix (FigureS1C), consistent with the recent reports that mitochondria might play a vital role in COVID-19. For instance, cytokine storm could induce iron dysfunction, such as hyperferritinemia, which can lead to the production of reactive oxygen species (ROS) and oxidative stress, and finally cause mitochondrial dysfunction, platelet damage, apoptosis. Abnormal platelets will cause clotting events and thrombus formation (Saleh et al., 2020). Howard Chang and college predicted that SARS-CoV-2 RNA enriched in host mitochondria (Wu et al., 2020), and mitochondria dysfunction was also identified in COVID-19 patients (Saleh et al., 2020).

Collectively, we identified HIS in SARS-CoV-2 genome, and the targeted human genome loci enriched with cytokines genes suggested that HIS may underly the clinical characteristics of COVID-19 patients and serve as a vital player in the pathological progression.

### Distribution and Conversation of HIS in 19 Species

SARS-CoV-2 belongs to the *Coronavirinae* subfamily, which contains six well-known human coronaviruses (HCoVs), including two alphacoronavirus (HCoV-299E, HCoV-NL63), two lineages A beta-coronavirus (HCoV-OC43, HCoV-HKU1), one lineage B beta-coronavirus (SARS-CoV), and one lineage C beta-coronavirus (MERS-CoV). To Figure out whether HIS is a common feature embedded across the HCoVs’ genome (The accession number for each HCoVs refers to TableS1), we analyzed the HCoVs and human genome sequence using the same criteria. Except for the five HIS in SARS-CoV-2 mentioned above, we overall identified 17 HIS, including five HIS in HCoV-HKU1, five HIS in HCoV-NL63, two HIS in HCoV-SARS, two HIS in HCoV-299E, two HIS in HCoV-OC43, one HIS in MERS-CoV (Figure2A). Like HIS-SARS-CoV-2 was abbreviated as HIS-SARS2, HIS-SARS-CoV and HIS-MERS-CoV were abbreviated as HIS-SARS and HIS-MERS, respectively (The detailed information of 22 HIS in HCoVs refers to TableS2.)

**Figure 2.**
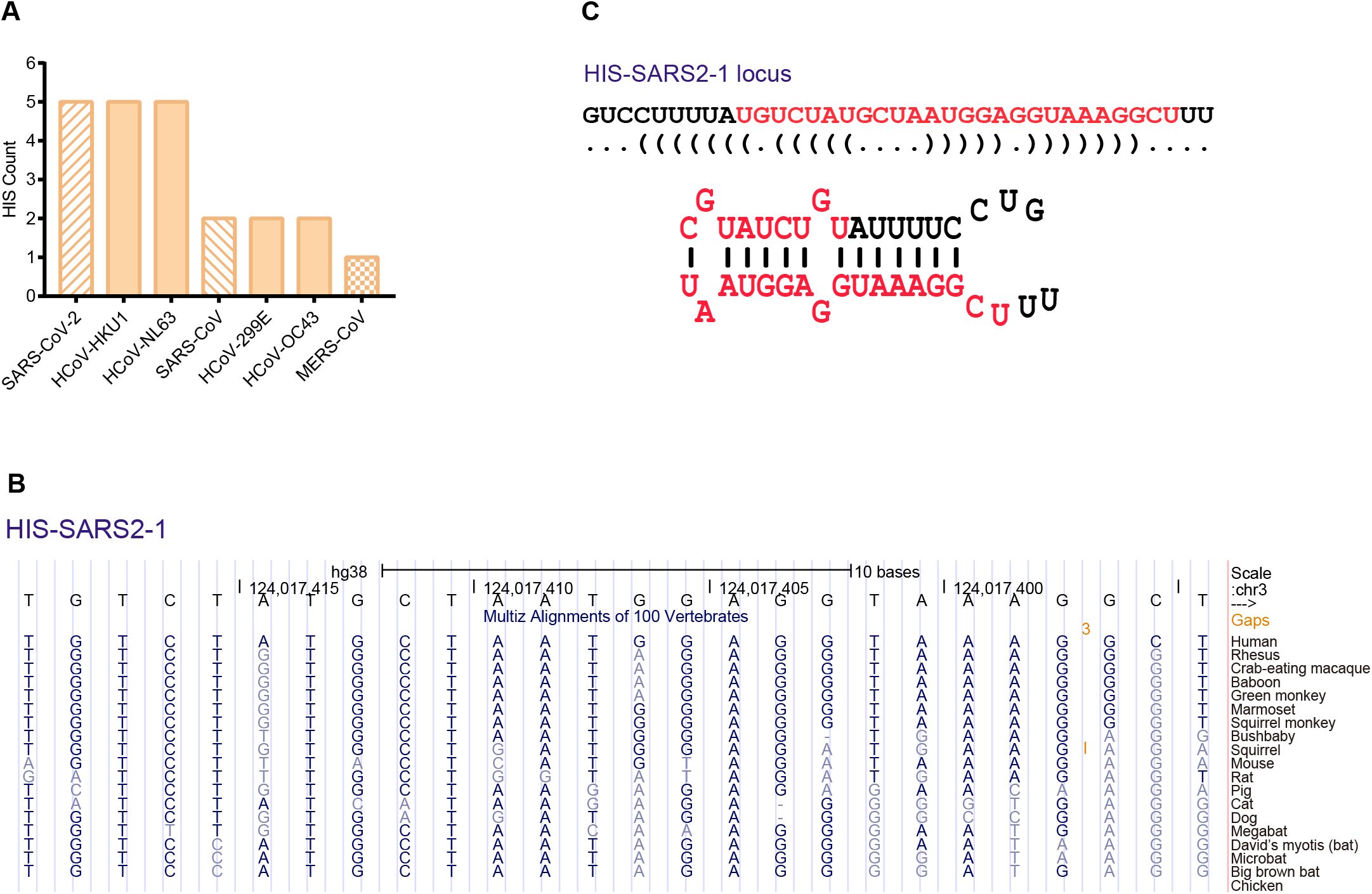
The Conservation of HIS across Species. **(A)** Number of HIS identified in seven known HCoVs. HIS, human identical sequences; HCoV, human coronavirus. **(B)** The sequence conservation of HIS-SARS2-1 in 19 species. The color of bases represents the conservation degree, dark blue represents the fully conserved, while light blue represents the mismatched base, single line (-) indicates no bases in the aligned species. In chicken, there doesn’t exist any conserved sequences. **(C)** The predicted RNA secondary structures of HIS-SARS2-1 and its adjunct bases (defined as pre-HIS-SARS2-1) by the algorithm of minimum free energy. The sequence in red represents HIS-SARS2-1 RNA.

These 22 HIS correspond to 35 identical sequences in human genome, which are distributed in 14 different chromosomes. It is noteworthy that one HIS may correspond to multiple loci in human genome, and chr6 (6/35) and chr4 (5/35) are the hotspots for HIS (FigureS2A), including five loci in MHC region of chr6 (TableS2). The length of HIS ranges from 24bp to 27 bp, while 18 out of 22 HIS are 24bp (TableS2). Meanwhile, the GC ratio range from 12.5% to 42.3%; however, GC ratio of 14 out of 22 (63.6%) HIS were no greater than 25% (FigureS2B). The universal distribution of HIS in HCoVs suggests a comprehensive role of these unheeded elements.

SARS-CoV-2 was reported to originate from bat (Zhou et al., 2020). However, many animals can be its potential host, including Malayan pangolins (Lam et al., 2020; Xiao et al., 2020), ferrets, and cats(Shi et al., 2020). We presumed that HIS might also exist in mediator hosts. To explore the distribution of HIS-SARS-CoV-2 across species, we analyzed the sequences of 19 species, including eight primates (human, rhesus, crab-eating macaque, baboon, green monkey, marmoset, squirrel monkey, bushbaby), squirrel, mouse, rat, pig, cat, dog, four bats (megabat, David’s myotis bat, microbat, big brown bat) and chicken (The accession number for each species mentioned here refers to TableS1). It was demonstrated clearly that all these five HIS-SARS2 were highly conserved in primates (Figure2B & FigureS3). Among them, HIS-SARS2-1, HIS-SARS2-4, and HIS-SARS2-5 were also highly conserved in bats and relatively less conserved in other mammals, such as squirrel, mouse, rat, cat, and dog (Figure2B, FigureS3C & FigureS3D); while HIS-SARS2-2 and HIS-SARS2-3 showed less conservation beyond primates (FigureS3B & FigureS3C). When it comes to the domesticated animals exampled by chicken, which is the main host for avian influenza, no conserved HIS were identified at all. The conservation of HIS across species hinted the propagation evolution pathway of SARS-CoV-2 and highlighted the unrevealed significance of HIS.

### HIS-SARS2 Forms miRNA-like Precursors

To investigate the potential functions of these HIS embedded fragments, we performed RNA secondary structures prediction (Mathews et al., 2004) and found that HIS-containing fragments were capable of forming the typical hairpin structure like miRNA precursors (Figure2C & FigureS4). Emerging evidence exhibits how human miRNAs interact with viral RNA, including direct binding (Chen et al., 2020c; Hosseini Rad Sm and McLellan, 2020; Nersisyan et al., 2020), thus it’s reasonable to suppose that virus-derived miRNA-like RNAs could also regulate human RNAs or even interact with human genome DNA elements. Indeed, SARS-CoV-encoded small viral RNAs (svRNAs) targeted 3′UTR of specific transcripts and repressed mRNA expression (Morales et al., 2017). Also, the function of small regulatory RNAs is largely dependent on their subcellular locations. It was predicted that SARS-CoV-2 RNA was highly enriched in the host nucleolus and mitochondria (Wu et al., 2020). Previously, our team revealed that miRNA could be allocated in the nucleus and target enhancer for gene activation (Xiao et al., 2017; Zou et al., 2017). Therefore, we suspected that HIS-SARS2 could interact with host enhancers and function like NamiRNA.

### HIS-SARS2 Activates Host Genes Involved in COVID-19 Pathogenic Processes

To investigate whether HIS can activate the host gene expression, we constructed vectors containing HIS fragments identified in SARS-CoV-2 and SARS-CoV. We transfected them into transformed human embryonic kidney cell HEK293T, human fetal lung fibroblast cell MRC5, and human umbilical vein endothelial cell (HUVEC), and detected the expression of surrounding as well as the distant genes. The expression level of coronavirus fragments was verified by qPCR (FigureS5A&B&C). *FBXO15*, *TIMM21*, and *CYB5A* lie the upstream of targeted locus of HIS-SARS2-4; among them, *FBXO15* was upregulated when HIS-SARS2-4 was transfected in HEK293T (Figure3A), MRC5 (Figure3B), and HUVEC (Figure3C); meanwhile, *TIMM21* and *CYB5A* upregulated in HEK293T (Figure3A) and MRC5 (Figure3B), respectively. As an E3 ligase subunit, *FBXO15* impairs mitochondrial integrity and induces lung injury in pneumonia (Chen et al., 2014). Both *TIMM21* (Mick et al., 2012) and *CYB5A* (Plitzko et al., 2013) encode the components of mitochondrial membrane, implying their fundamental roles in mitochondria, the dysfunction of which is tightly correlated with COVID-19 pathogenesis (Saleh et al., 2020). Similarly, HIS-SARS2-1 overexpression upregulated the upstream gene *KALRN* in HEK293T cell (FigureS5D). *KALRN* plays a significant role in the development of sarcoidosis (Besnard et al., 2018), a systemic inflammatory disease involved in multiple organs throughout the body, including the kidney and lungs, implying *KALRN* may get involved in inflammatory response after SARS-CoV-2 infection. Except for the adjacent genes, HIS can also upregulate distant genes. The transfected HIS-SARS2-3 upregulated *MYL9* in HEK293T and *MRC5* (Figure3D) and *EPN1* (Figure3E) in MRC5 and HUVEC (Figure3F). *MYL9* was demonstrated to be a functional ligand for CD69 that regulate airway inflammation (Hayashizaki et al., 2016). These data suggest HIS contribute to COVID-19 pathogenesis by upregulating the adjacent or distant genes, which may cause mitochondrial dysfunction or participate in the inflammatory response.

**Figure 3.**
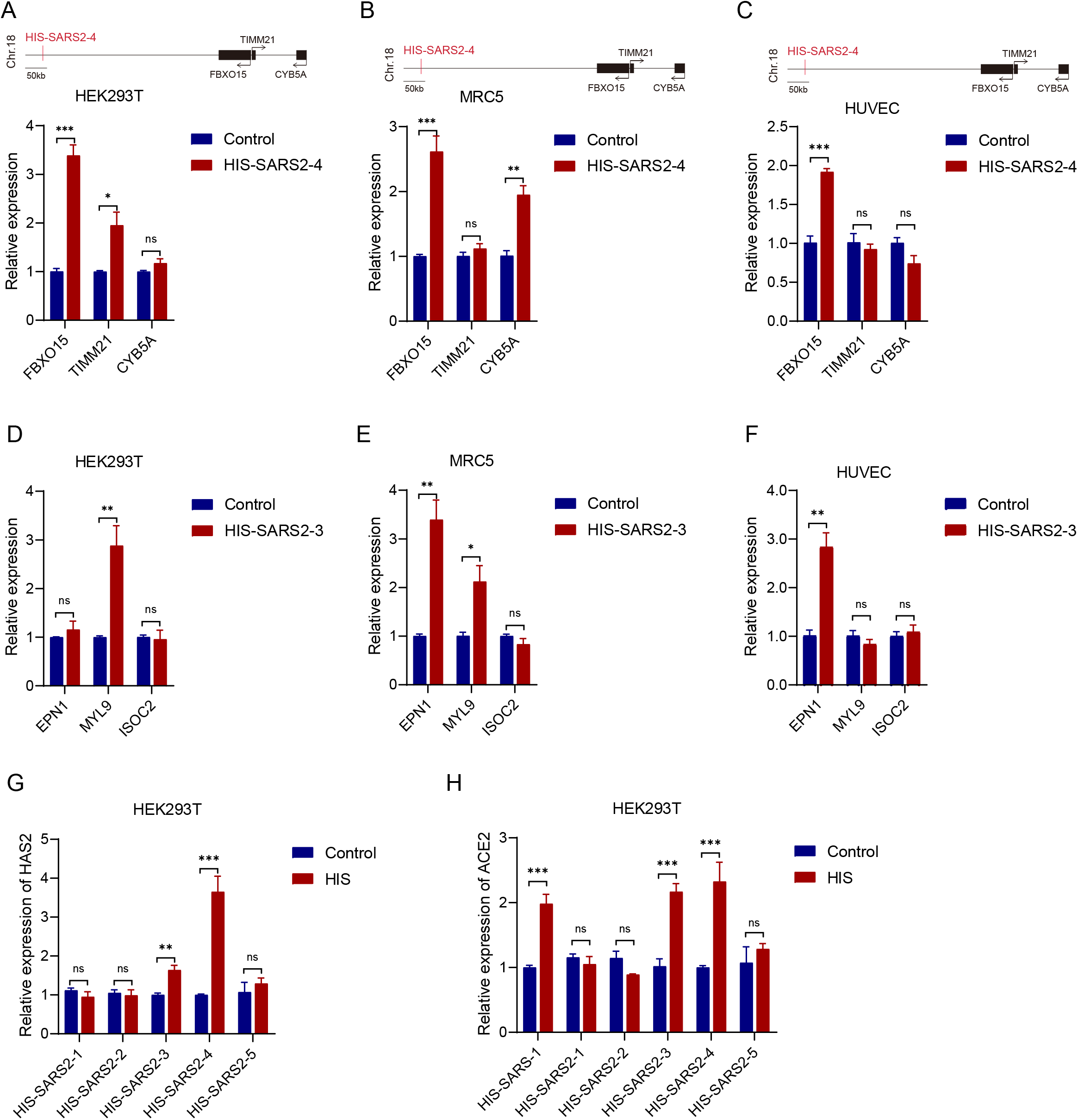
HIS-SARS2 Activates Genes Related to COVID-19 Pathology. **(A)(B)(C)** The relative mRNA expression of the neighboring genes *FBXO15*, *TIMM21*, and *CYB5A* after HIS-SARS2-4 vector transfected in HEK293T cells (A), MRC5 cells (B), and HUVEC cells (C). **(D)(E)(F)** The relative mRNA expression of the distant gene *EPN1*, *MYL9*, and *ISOC2* after HIS-SARS2-3 vector transfected in HEK293T cells (D), MRC5 cells (E), and HUVEC cells (F). **(G)** The relative mRNA expression of hyaluronan synthase *HSA2* after five HIS-SARS2 vectors transfected in HEK293T cells. HIS-SARS2-3 and HIS-SARS2-4 upregulates the expression of *HSA2*. **(H)** The relative mRNA expression of *ACE2*, which encodes the cell entry receptor of SARS-CoV-2, after HIS-SARS-1 and five HIS-SARS2 vectors transfected in HEK293T cells. HIS-SARS-1, HIS-SARS2-3, and HIS-SARS2-4 upregulates the expression of *ACE2*. In all Figure s above, the *y*-axis indicates the mRNA fold changes to *GAPDH* detected by RT-qPCR. *P*-values were calculated using the unpaired, two-tailed Student’s t test by GraphPad Prism 7.0. *, *P*<0.05; **, *P*<0.01; ***, *P*<0.001; ns, not significant.

Like SARS-CoV-2, SARS-CoV is a typical fetal beta-coronavirus. Patients who contracted these two largely share similar symptoms (Chen et al., 2020a). Transfection of HIS-SARS-1 could also activate neighboring genes, such as *HAS2*, *ZHX2*, and *DERL1* in HEK293T (FigureS5E), MRC5 (FigureS5F), and HUVEC (FigureS5G). Importantly, *HAS2* encodes the critical enzyme for the production of hyaluronan, which is a kind of glycosaminoglycan that plays a fundamental role in the inflammatory response. The accumulation of hyaluronan in the lung has been recognized in ARDS for more than three decades (Hallgren et al., 1989). Besides, ARDS is the most typical clinical manifestation of severe COVID-19 cases (Guan et al., 2020; Wang et al., 2020a). In that, it’s natural to testify whether HIS-SARS2 could activate *HAS2*. Surprisingly, we found HIS-SARS2-3 and HIS-SARS2-4 could activate the expression of *HAS2* about 2 ~4 folds in HEK293T (Figure3G) and MRC5 (FigureS5H); however, the expression of *HAS2* was upregulated more than ten folds in HUVEC (FigureS5I), suggesting that small vascular cell may be the main target of SARS-COV2 infection. In this case, HIS might largely explain most clinical symptoms of COVID-19 patients, including ARDS and microvessels injury, by upregulating *HAS2* and the level of hyaluronan, which receives very limited attention in COVID-19 progression.

Additionally, we noticed that *ACE2* could also be upregulated by HIS-SARS2-3 and HIS-SARS2-4 (Figure3H). ACE2 is the essential receptor for both SARS-CoV-2 and SRAS-CoV to enter the host cell (Hoffmann et al., 2020; Zhou et al., 2020). These data indicate HIS-SARS2 not only induce the clinical manifestations of COVID-19 patients but also enhance the spreading of SARS-CoV-2 by promoting the level of ACE2 and making it more susceptible, which partially explain the global threat of pandemic for COVID-19.

### HIS-SARS2 Activates the Host Genes through NamiRNA-Enhancer Network

To explore the underlying mechanism of HIS regulated gene expression. We transfected HEK293T with HIS fragments and performed H3K27ac ChIP-qPCR. It clearly showed that HIS fragments could induce the enrichment of H3K27ac at HIS targeted loci (Figure4A). H3K27ac marks enhancer or super-enhancer, which is critical for gene activation. This indicates that HIS regulated gene activation is largely dependent on activating the targeted enhancers. JQ1 is a potent inhibitor of the BET family proteins, which functions by binding to H3K27ac. After the treatment of JQ1, the upregulation of HIS targeted genes was abolished (Figure4B&C&D), which was also the case in HIS-SARS-1 (FigureS6A), further supporting that HIS mediated gene regulation is achieved through enhancer. To further confirm whether HIS were indispensable for gene activation, we knocked down the HIS expression by Cas13d and found the activation of targeted genes was abolished (Figure4E&F&G), which was also the case in HIS-SARS-1 (FigureS6B), indicating the specificity of HIS for regulating host gene. The behaviors of HIS-SARS2 RNA are highly similar to our previously identified NamiRNAs, which activate genes as enhancer triggers in the nucleus (Xiao et al., 2017; Zou et al., 2017). Overall, these data suggest that HIS activates the host gene through the NamiRNA-enhancer network.

**Figure 4.**
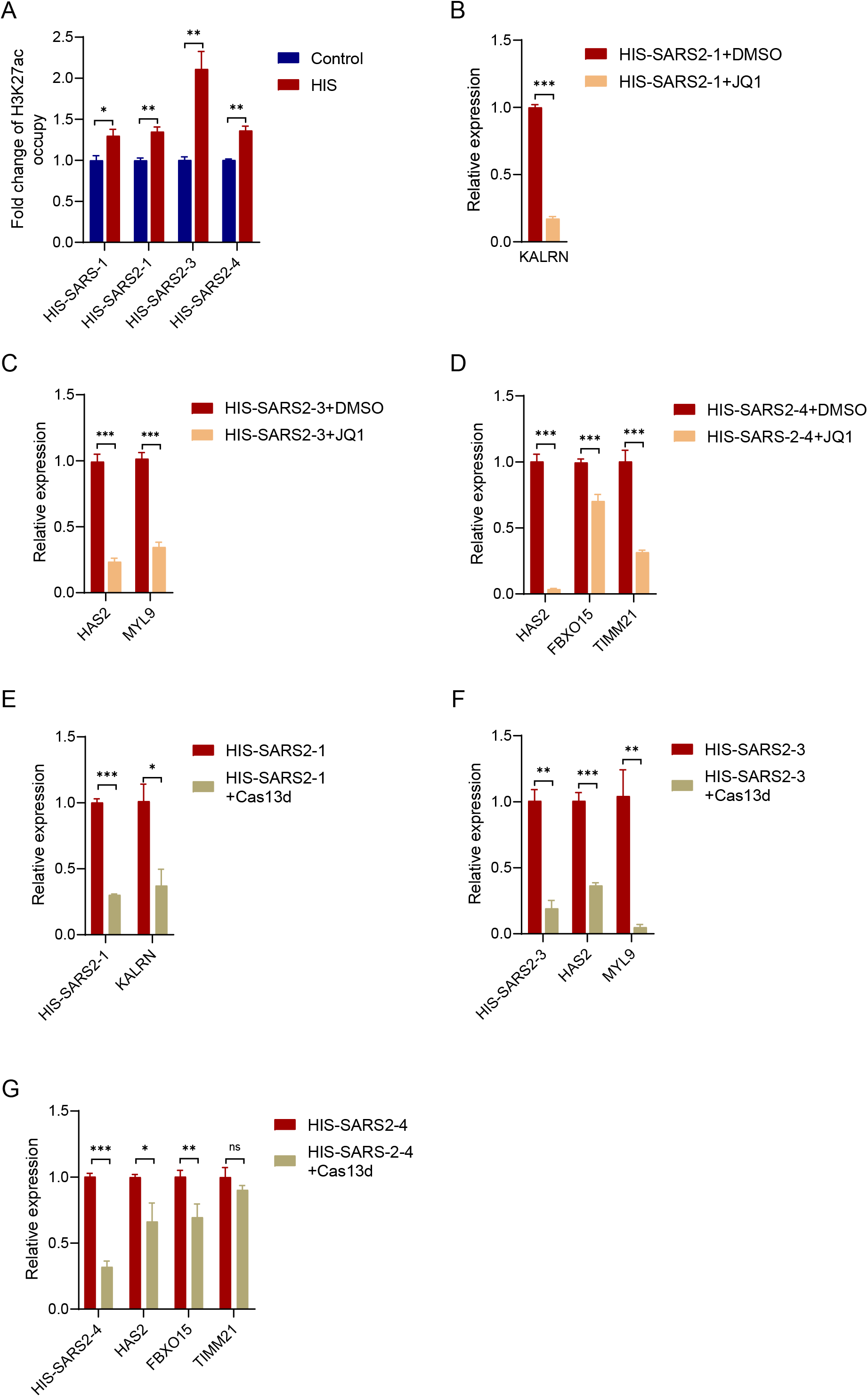
H3K27ac and HIS are Essential for Upregulation of HIS Targeted Genes. **(A)** The enrichment of enhancer marker H3K27ac after HIS-SARS-1, HIS-SARS2-1, HIS-SARS2-3, and HIS-SARS2-4 transfection. The *y*-axis indicates the DNA which was pulled down by H3K27ac ChIP and detected by qPCR. **(B)** The relative mRNA expression of HIS-SARS2-1 targeted gene *KALRN* in HEK293T cells, which were transfected with HIS-SARS2-1 precursor vector and then treated with 500 nM JQ1 for 24 h. JQ1 abolishes the upregulation of HIS-SARS2-1 targeted gene *KALRN*. **(C)** The relative mRNA expression of HIS-SARS2-3 targeted genes *HAS2* and *MYL9* in HEK293T cells, which were transfected with HIS-SARS2-3 precursor vector and then treated with 500 nM JQ1 for 24 h. JQ1 abolishes the upregulation of HIS-SARS2-3 targeted gene *HAS2* and *MYL9*. **(D)** The relative mRNA expression of HIS-SARS2-4 targeted genes *HAS2*, *FBXO15*, and *TIMM21* in HEK293T cells, which were transfected with HIS-SARS2-4 precursor vector and then treated with 500 nM JQ1 for 24 h. JQ1 abolishes the upregulation of HIS-SARS2-4 targeted gene *HAS2*, *FBXO15*, and *TIMM21*. **(E)** The relative mRNA expression of HIS-SARS2-1 targeted gene *KALRN* in HEK293T cells, which were transfected with HIS-SARS2-1 precursor vector and then knocked down by Cas13d. Knockdown HIS-SARS2-1 by Cas13d abolished the upregulation of HIS-SARS2-1 targeted gene *KALRN*. **(F)** The relative mRNA expression of HIS-SARS2-3 targeted genes *HAS2* and *MYL9* in HEK293T cells, which were transfected with HIS-SARS2-3 precursor vector and then knocked down by Cas13d. Knockdown HIS-SARS2-3 by Cas13d abolished the upregulation of HIS-SARS2-3 targeted genes *HAS2* and *MYL9*. **(G)** The relative mRNA expression of HIS-SARS2-4 targeted genes *HAS2*, *FBXO15*, and *TIMM21* in HEK293T cells, which were transfected with HIS-SARS2-3 precursor vector and then knocked down by Cas13d. Knockdown HIS-SARS2-4 by Cas13d abolished the upregulation of HIS-SARS2-4 targeted genes *HAS2*, *FBXO15*, and *TIMM21*. In (B)(C)(D)(E)(F)(G), The *y*-axis indicates the relative mRNA fold changes to *GAPDH* detected by RT-qPCR. In all Figure s, *P*-values were calculated using the unpaired, two-tailed Student’s t test by GraphPad Prism 7.0. *, *P*<0.05; **, *P*<0.01; ***, *P*<0.001; ns, not significant.

### HIS-SARS2 RNA Binds to Identical Human Loci Stabilized by AGO2

To verify the binding between HIS and its targeted sequences in human genome, we synthesized HIS-SARS2 and their target DNA fragments (TableS3). We detected the interaction between HIS-SARS2-1 and its target DNA by UV absorption spectra. Generally, after the hybridization, the absorbance of the hybrid strand will decrease. When HIS-SARS2-1 RNA and its target ssDNA (HIS-SARS2-1 DNA-S, S represents sense strand) were hybridized, the absorbance significantly decreased compared with the theoretically calculated and the mixed (before hybridization) group, indicating that HIS-SARS2-1 can hybridize its target DNA and form double strands (Figure5A). Comparing the relative absorbance change, we found when 50% product changed, the corresponding T_m_ value for HIS-SARS2-1-RNA-DNA hybrid (62.7°C) was higher than dsDNA (60.3°C) (Figure5B), the increase in which reflected the robustness of HIS-SARS2-1-RNA-DNA hybrid over dsDNA.

**Figure 5.**
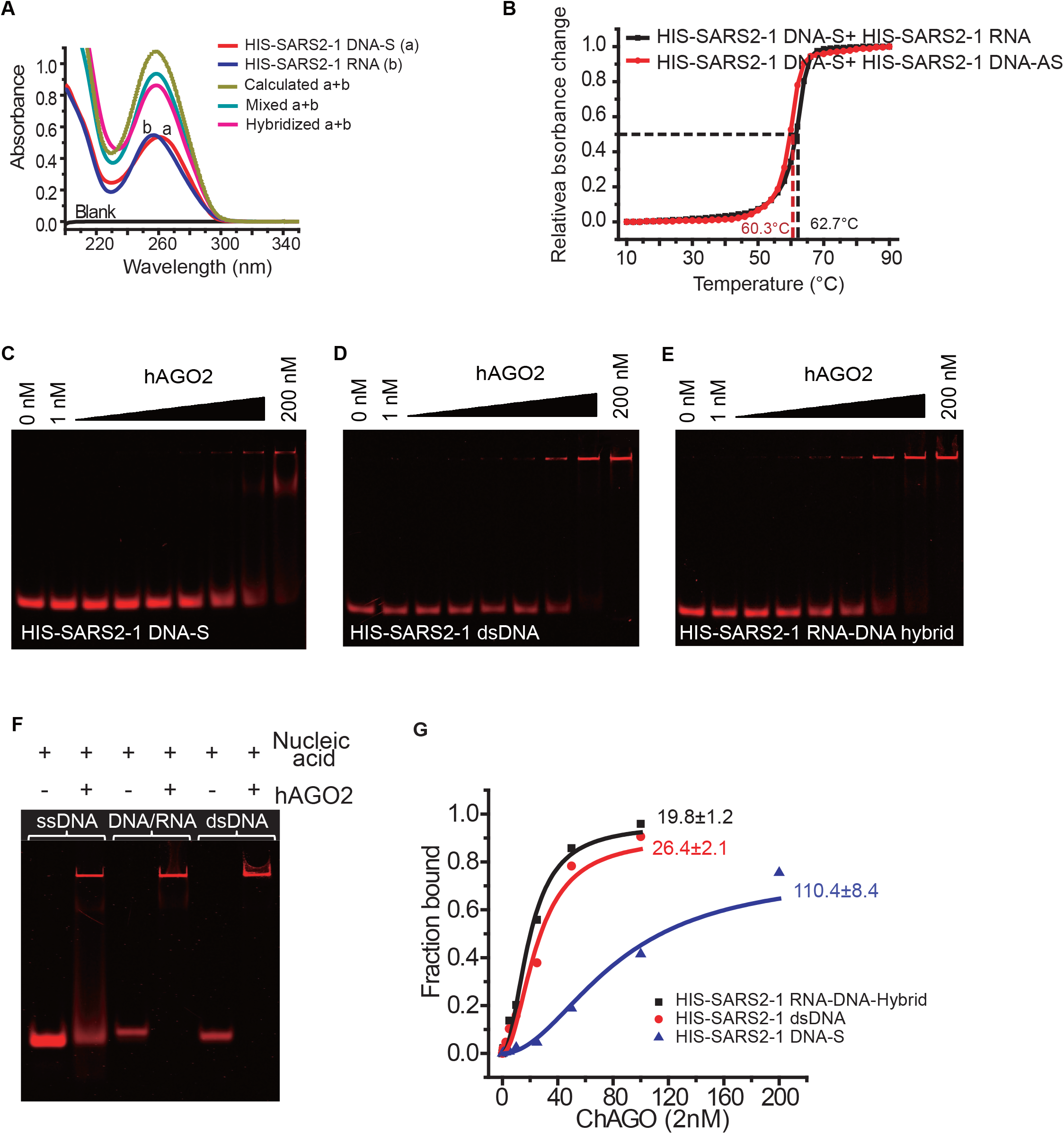
HIS-SARS2 RNA Binds to Homologous Sequence of Human Genome for Gene Activation. **(A)** The UV absorption spectra of 2 μM HIS-SARS2-1 DNA (red curve a), 2 μM HIS-SARS2-1 RNA (blue curve b), mixed HIS-SARS2-1 DNA and RNA (dark cyan curve), hybridized mixture of HIS-SARS2-1 (magenta curve), and the simple sum of curve a and b (dark yellow dotted curve). **(B)** The relative absorbance change of 2 μM hybridized mixture of HIS-SARS2-1 and the complementary RNA-HIS-SARS2-1(black square) or HIS-SARS2-1 and the complementary ssDNA (red circle) at 260 nm versus temperature. **(C)** Gel showing of 25 nM 5’Cy5-HIS-SARS2-1 in the presence of increasing concentrations of hAGO2. **(D)** Gel showing of 25 nM 5’Cy5-HIS-SARS2-1 DNA duplex in the presence of increasing concentrations of hAGO2. **(E)** Gel showing of 25 nM 5’Cy5-HIS-SARS2-1 DNA-RNA hybrid in the presence of increasing concentrations of hAGO2. **(F)** Gel showing 25nM 5’Cy5-HIS-SARS2-1 (lane 1, 2), 5’Cy5-HIS-SARS2-1 DNA-RNA hybrid (lane 3, 4), 5’Cy5-HIS-SARS2-1 DNA duplex (lane 5, 6) in the absence and presence of 100 nM hAGO2. **(G)** The bound fraction of 5’Cy5-HIS-SARS2-1 (blue triangle), 5’Cy5-HIS-SARS2-1 DNA-RNA hybrid (black square) and 5’Cy5-HIS-SARS2-1 DNA duplex (red circle) probes in the presence of different concentration of hAGO2.

AGO2 is the key factor for miRNA and RNAi in function. Previously, we have shown that AGO2 was also essential for NamiRNA-enhancer-gene activation pathway (Liang et al., 2019). Here we interrogated whether AGO2 was crucial for HIS-induced gene activation. We accessed the interaction between human AGO2 (hAGO2) and HIS-SARS2-1 DNA-S, HIS-SARS2-1 dsDNA, HIS-SARS2-1 RNA-DNA-Hybrid by electrophoretic mobility shift assay (EMSA) assay. It showed that hAGO2 could partially bind to HIS-SARS2-1 DNA-S in high hAGO2 concentration (Figure5C), while the interaction between hAGO2 and HIS-SARS2-1 dsDNA or HIS-SARS2-1 RNA-DNA-Hybrid was more intense (Figure5D&E), which was verified in a fixed concentration of hAGO2 (Figure5F). We simulated the hAGO2 binding curves of these three forms of nucleic acid and calculated the corresponding dissociation constant (K_d_). It clearly demonstrated that HIS-SARS2-1 RNA-DNA-Hybrid has the smallest K_d_, indicating the strongest binding ability. Comparing with HIS-SARS2-1 RNA-DNA-Hybrid, HIS-SARS2-1 dsDNA exhibited a relatively low binding capability, and HIS-SARS2-1 DNA-S bound weakly to hAGO2 (Figure5G).

These data highly suggested that HIS-SARS2 RNAs could bind to their targeted loci in the host, exampled typically here by the identical sequences in human genome, and activate genes related to COVID-19 pathology via AGO2-dependent NamiRNA-enhancer network.

### Hyaluronan Acts as a Predictor of Progression and Potential Therapeutic Target for COVID-19 Treatment

SARS-CoV and SARS-CoV-2 are similar in many aspects; especially, the infected patients are characterized by ARDS and pulmonary fibrosis. It has been reported that SARS-CoV infection promotes acute lung injury by regulating hyaluronan level, which is mediated by *HAS3* (Hu et al., 2012). Among HIS-activated genes, *HAS2* arose our special attention for its ability to regulate hyaluronan level. Accordingly, we found that HIS-SARS-1, HIS-SARS2-3, and HIS-SARS2-4 significantly upregulated the hyaluronan level in supernatant of culture medium for HEK293T (Figure6A), which was also the case in MRC5 (FigureS7A). Therefore, we speculated that hyaluronan may get involved in COVID-19 pathogenesis and progression.

**Figure 6.**
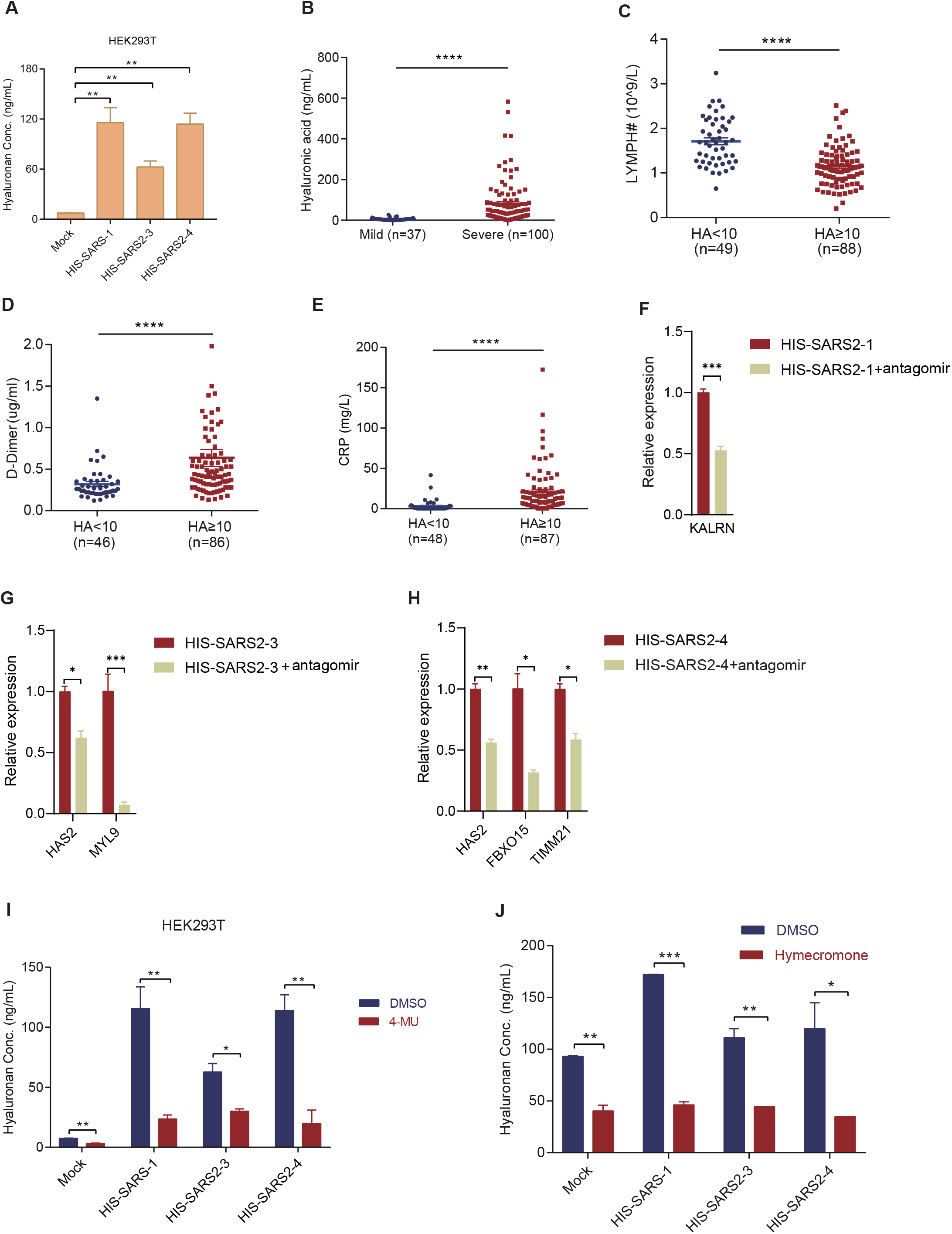
Hyaluronan as a Potential Therapeutic Target of COVID-19. **(A)** Hyaluronan released in cell culture supernatants of Mock, HIS-SARS-1, HIS-SARS2-3, and HIS-SARS2-4 overexpressed in HEK293T cell. *P*-values were calculated using the unpaired, two-tailed Student’s t test. **, *P*<0.01. **(B)** Hyaluronan levels in mild patients (n=37) and severe patients (n=100). The *y*-axis indicates the hyaluronan levels detected by ELISA. *P*-values were calculated using the two-tailed nonparametric Mann–Whitney test by GraphPad Prism 7.0. ****, *P*<0.0001. **(C)** Hyaluronan level is negatively correlated with lymphocytes number in patients’ plasma. HA (hyaluronic acid) level (10 ng/mL) functions as a discriminator for patients’ lymphocytes number. For patients whose hyaluronan <10 ng/mL (49 patients), the mean value is 1.71×10^9^/L; while for patients whose hyaluronan ≥10 ng/mL (88 patients), the mean value is 1.15×10^9^/L. *P*-values were calculated using the two-tailed nonparametric Mann–Whitney test by GraphPad Prism 7.0. ****, *P*<0.0001. **(D)** Hyaluronan level is positively correlated with D-Dimer level in patients’ plasma. HA (hyaluronic acid) level (10ng/mL) functions as a discriminator for patients’ D-Dimer levels. For patients whose HA<10ng/mL (46 patients), the mean value is 0.32 μg/ml; while for patients whose HA≥10 ng/mL (86 patients), the mean value is 0.64 μg/ml. *P*-values were calculated using the two-tailed nonparametric Mann–Whitney test by GraphPad Prism 7.0. ****, *P*<0.0001. **(E)** Hyaluronan level is positively correlated with C-reactive protein (CRP) level in patients’ plasma. HA (hyaluronic acid) level (10ng/mL) functions as a discriminator for patients’ CRP levels. For patients whose HA<10ng/mL (48 patients), the mean value is 3.18 mg/L; while for patients whose HA≥10ng/mL (87 patients), the mean value is 20.75 mg/L. *P*-values were calculated using the two-tailed nonparametric Mann–Whitney test by GraphPad Prism 7.0. ****, *P*<0.0001. **(F, G, H)** Antagomirs of HIS-SARS2-1, HIS-SARS2-3, and HIS-SARS2-4 block the upregulation of the targeted gene *KALRN* (F), *MYL9* (G), *HAS2*, *FBXO15*, and *TIMM21* (H), respectively. *P*-values were calculated using the unpaired, two-tailed Student’s t test by GraphPad Prism 7.0. *, *P*<0.05; **, *P*<0.01; ***, *P*<0.001; ****, *P*<0.0001. **(I)** HA released in cell culture supernatants of Mock, HIS-SARS-1, HIS-SARS2-3, or HIS-SARS2-4 overexpressed in HEK293T after treated with 100 μM 4-MU and DMSO as control. *P*-values were calculated using the unpaired, two-tailed Student’s t test. *, *P*<0.05; **, *P*<0.01. **(J)** HA released in cell culture supernatants of Mock, HIS-SARS-1, HIS-SARS2-3, and HIS-SARS2-4 overexpressed in HEK293T after treated with 200 μg/ml hymecromone and DMSO as control. *P*-values were calculated using the unpaired, two-tailed Student’s t test. *, *P*<0.05; **, *P*<0.01; ***, *P*<0.001.

To further decipher the fundamental role of hyaluronan in COVID-19, we collected the plasma of COVID-19 patients who have been hospitalized at The Shanghai Public Health Clinical Center. We categorized patients into mild (n=37) and severe (n=100) groups based on the characteristic pneumonia features of chest CT. In severe patients, the mean value of hyaluronan (80.39ng/mL) (Figure6B) was significantly higher than that in mild patients (5.70ng/mL), which was supported by the recent report that hyaluronan level was higher in severe patients (Ding et al., 2020). Hyaluronan can encompass a large volume of water, which enable it to determine the water content in specific tissues (Turino and Cantor, 2003). It has been reported that the extravascular lung water volume is positively correlated with hyaluronan level in normal animal lungs (Bhattacharya et al., 1989). In that, the water absorption characteristics of hyaluronan rationalize the possibility that increased hyaluronan, which was induced by the upregulated *HAS2* in lung cells after SARS-CoV-2 infection, bind lots of water and form jelly-like substances, underlying the ground-glass opacity commonly occurred in COVID-19 patients. Besides, higher level of hyaluronan in severe patients’ plasma suggests hyaluronan may act as a predictor of COVID-19 progression, especially when physicians making a quick decision in the clinic for patients who will need critical medical care.

To test whether hyaluronan level alone could be an indicator for characterizing the clinical manifestations of COVID-19 patients, we classified patients into a mild and severe group based on the hyaluronan level. We found that in severe patients, the level of lymphocytes decreased (Figure6C), D-Dimer increased (Figure6D), and C-reactive protein (CRP) upregulated (Figure6E), which was supported by the report that compared non-intensive care unit (ICU) patients, D-dimer was significantly upregulated, while lymphocyte count decreased (Wang et al., 2020a). Additional evidence showed that lymphocyte count, C-reactive protein, and D-dimer were significantly associated with COVID-19 severity (Wang et al., 2020b). These data suggest hyaluronan alone can function as a potential indicator for COVID-19 severity.

Based on the discoveries that HIS regulated *HAS2* and hyaluronan level contributed to the clinical features of COVID-19 patients, we speculate the downregulation of hyaluronan by HIS antagomirs might be an effective way to abolish the upregulation of inflammatory genes and even relieve the clinical symptoms. To testify this strategy, we transfected cells with antagomirs and found antagomirs of HIS-SARS2-1, HIS-SARS2-3, and HIS-SARS2-4 downregulated the inflammatory genes which had been upregulated, such as *KALRN* (Figure6F), *HAS2*, *MYL9* (Figure6G), *FBXO15*, and *TIMM21* (Figure6H). Antagomir of HIS-SARS-1 also downregulated the *HAS2* and *ZHX2* (FigureS7B). These data reveal the druggable potential of HIS antagomir for COVID-19 treatment.

To further test whether hyaluronan could function as a target for COVID-19 progression inhibition, we treated HEK293T with 4-Methylumbelliferone (4-MU), a hyaluronan synthesis inhibitor (Nagy et al., 2015). After treatment, hyaluronan was downregulated in all groups (Figure6I), which was also the case in MRC5 (FigureS7C). We noticed there was a prescription drug, hymecromone, which also function as a hyaluronan inhibitor. Similarly, we treated HEK293T with DMSO-dissolved hymecromone and found hyaluronan reduced accordingly in the cell culture supernatants (Figure6J).

Taken together, hyaluronan promoted by HIS is emerging as a novel target for COVID-19 treatment, and the inhibition of *HAS2* by HIS antagomir or 4-Methylumbelliferone might be novel strategies to block the progression of COVID-19.

## Discussion

It is of critical need to understand the mechanism of how SARS-CoV-2 causes inflammation cytokines storm in the host response. In this current study, we illustrated that human identical sequences (HIS) of SARS-CoV-2 activate host cytokines through the NamiRNA-enhancer-gene activation network. Ectopic expression of these fragments containing HIS-SARS2 promotes H3K27ac enrichment at their corresponding regions in host cells. It is noteworthy that these HIS of SARS-CoV-2 can bind steadily to their corresponding regions of human DNA fragments *in vitro*. Importantly, HIS activate *HAS2* expression and increase hyaluronan level, while COVID-19 patients who have higher plasma hyaluronan levels tend to show severe symptoms. These findings contribute to understanding the progression of COVID-19 after SARS-CoV-2 infection and may provide novel therapeutic targets.

A growing number of studies indicate that the infection of various viruses, including both DNA and RNA viruses, has species-specific and organ-specific signatures (De Meyer et al., 1997; Iwamoto et al., 2004; Ohlund et al., 2019; Rothenburg and Brennan, 2020). For example, humans and chimpanzees are the known unique hosts to be infected naturally by Hepatitis C virus (HCV) (Sandmann and Ploss, 2013). Notably, the liver is the principal site of HCV infection. The pathogenesis of HCV infection-causing progressive liver diseases is thought to be an uncontrolled inflammatory response within the change of inflammatory cytokines (Li et al., 2018). Of note, not all the infection of viruses is virulent for their hosts. One of the typical examples is that human immunodeficiency virus type 1 (HIV-1) is pathogenic only for human, and does not affect the health of the primary host (Nomaguchi et al., 2012). Similarly, SARS-CoV-2 infection in human results in COVID-19 without being fatal for its other potential hosts, including bats and pangolins (Xiao et al., 2020; Zhou et al., 2020). By comparing different primates with SARS-CoV-2 genome, we found five conserved fragments with high similarity in human. Interestingly, the conserved fragments were discovered between SARS-CoV-2 and their hosts, including bats. Moreover, we also identified HIS between other highly pathogenic beta-coronaviruses (such as SARS-CoV and MERS-CoV) and different primates. Consistent with these points, HIS was found within other pathogenic viruses, including avian influenza virus, swine flu virus, rabies virus, coxsackievirus, influenza A virus, HIV, Ebolavirus, and Zika virus (TableS4). Therefore, it is a universal phenomenon that the conserved fragments, which were termed here as "host identical sequences (HIS)", are ubiquitous between viruses and their hosts, implying HIS could be the determinants of viral susceptibility and pathogenicity. In this case, it is natural to speculate that HIS from the pathogen genome may help to trace the lines of the host, especially for the mediated host during SARS-CoV-2 virus evolution. Since HIS activate host genes through enhancer and enhancer is well known for its tissue-specific signature, it is paramount to investigate the virus-host cell interaction and to understand why viruses usually show organ or tissue-specific pathogenesis.

In recent years, multiple DNA and RNA viruses have been demonstrated to produce miRNA-like non-coding RNAs (Mishra et al., 2019). Surprisingly, bioinformatics analysis revealed that the fragments containing HIS in SARS-CoV-2 could produce the primary structure of precursor miRNA (pre-miRNA). We have previously clarified that miRNAs can activate genome-wide gene transcription by targeting enhancers (Xiao et al., 2017). Excitingly, further bioinformatics analysis (Betel et al., 2010) indicated that enhancers of lung targeted by these HIS could regulate 298 genes (TableS5) overlapping with recently published transcriptome sequencing data of the bronchoalveolar lavage fluid (BALF) in COVID-19 patients (Xiong et al., 2020), implying HIS probably play a key role during the progression of COVID-19. An increasing number of researches point to SARS-CoV-2 infection as causing multiple organ damage involved in the lung, kidney, and liver by activating inflammation response (Wiersinga et al., 2020). Additionally, the fibrinolytic system in COVID-19 patients is also destroyed by SARS-CoV-2 infection (Levi et al., 2020). To confirm whether HIS affect inflammation in these organs, different fragments of HIS were transfected in selected human cell lines, including HEK293T, MRC5, HUVEC. Interestingly, HIS-SARS2-1 can upregulate the expression of its upstream *KALRN* in HEK293T, which contributes to the development of sarcoidosis (Besnard et al., 2018), causing a systemic inflammatory in multiple organs such as kidney and lungs. Alternatively, the adjacent gene *FBXO15* and the distant gene *MYL9* were increased by transfected HIS-SARS2-3 and HIS-SARS2-4 in HEK293T. Of note, both *FBXO15* and *MYL9* are closely related to inflammation response (Chen et al., 2014; Hayashizaki et al., 2016). Similarly, HIS-SARS2-4 promoted the expression of its upstream gene *CYB5A* and *FBXO15* in MRC5. Consistent with these results, HIS in SARS-CoV also activate neighboring genes, such as *HAS2*, *ZHX2*, and *IGF2R*. Collectively, all the above evidence emphasizes that HIS can indeed cause inflammation response by activating gene expression, supporting this novel mechanism underlying the viral pathogenicity.

miRNAs can activate gene transcription epigenetically by increasing H3K27ac enrichment at their target enhancers in our previous study (Xiao et al., 2017). Interestingly, ChIP-qPCR confirmed that H3K27ac was enriched in the HIS-SARS2-4 region of the human genome by the corresponding HIS fragment in HEK293T. Then we treated HIS-SARS2-4 HEK293T cell with JQ1, which is an inhibitor of BRD4, resulting in the preferential loss of enhancer (Loven et al., 2013). Meanwhile, blocking H3K27ac with JQ1 remarkably downregulated *FBXO15* expression in HEK293T, suggesting enhancer is essential for HIS mediated gene activation. To investigate the specificity of this activation process, we designed the antagomir against HIS-SARS2-4 and found that *FBXO15* was decreased in HIS-SARS2-4 transfected HEK293T cell, which was further confirmed by degrading the HIS-SARS2-4 with CasRx system for efficiently and functionally knocking down targeted genes (He et al., 2020). As AGO2 may serve as a guide mediating the binding of miRNAs to their enhancers, resulting in gene activation (Liang et al., 2019), we found that AGO2 was also involved in HIS induced host gene activation. Clearly, HIS-SARS2-4 RNA could hybridize its target ssDNA and form stable double strands, which can be stabilized by hAGO2 by reducing dissociation constant (K_d_), supporting that AGO2 guiding the binding of miRNA on their target enhances. These results demonstrate that HIS can activate gene transcription epigenetically, and inhibition or degradation of HIS may act as a new strategy for controlling the virus-induced disease.

Importantly, we identified that hyaluronan could be a novel target for COVID-19 treatment. At first, *HAS2* located on the upstream of HIS targeted site in the human genome attracted our interest, the major enzyme responsible for hyaluronan synthesis (Csoka et al., 2001). Accumulation of hyaluronan is closely associated with ARDS (Hallgren et al., 1989), the typical symptom in patients of SARS-CoV, SARS-CoV-2, and MERS-CoV infection. As expected, HIS-SARS-CoV and HIS-SARS2 can activate dramatically *HAS2* expression in host cells, causing the upregulation of hyaluronan in cell medium supernatant. Clinically, hyaluronan is significantly increasing in severe COVID-19 patients with ground-glass opacity by the chest CT scan, and the level of hyaluronan is correlated with the clinical prognosis of patients with COVID-19 (Ding et al., 2020). Interestingly, decreased lymphocytes, increased D-Dimer and C-reactive protein show up more often in severe patients compared to mild patients distinguished by their hyaluronan level, which is in surprising unanimity with the pathological features of ICU COVID-19 patients (Wang et al., 2020a). In other words, hyaluronan level alone could be an indicator for characterizing the clinical progression of COVID-19 patients. Notably, the total number of T cells reduced significantly in COVID-19 patients compared to the normal levels (Qin et al., 2020). Such discrepancies may be owing to the binding of hyaluronan and its ligand CD44, which can induce the death of activated T cells (McKallip et al., 2002). Another ligand HABP2 (also called factor VII-activating protease), which conjugate with hyaluronan, plays an important role in blood coagulation by activating the pro-urokinase-type plasminogen activator (Kanse et al., 2008), which may cause the dysregulation of the fibrinolytic system in COVID-19 patients. In addition, HABP2 aggravates the disruption of the hyaluronan-mediated endothelial cell barrier (Mambetsariev et al., 2010), which may explain well the sudden brain hemorrhage of ICU patients with COVID-19 (Barrios-Lopez et al., 2020). Further studies will be performed to elucidate the latent mechanism of hyaluronan causing these clinical characteristics during the progression of COVID-19. 4-Methylumbelliferone is reported to be an efficient inhibitor for hyaluronan production (Nagy et al., 2015). 4-MU treatment can reverse the upregulation of hyaluronan in response to HIS in HEK293T and MRC5. Additionally, hyaluronan level was also decreased in HEK293T after the treatment of hymecromone, which is one of the commercially available hyaluronan inhibitor drugs for cholecystitis, cholelithiasis, and cholecystectomy syndrome in China. Overall, decreasing hyaluronan could improve most of the clinical symptoms of COVID-19 patients and serve as a promising therapeutic target for SARS-CoV-2 and other related diseases.

Taken together, our results presented here indicate a novel mechanism that SARS-CoV-2 can induce host response by regulating host gene expression through the direct interaction between their vsRNAs and chromatin enhancers in host during COVID-19 progression. Given these findings, blocking the HIS with nucleic acid drugs or inhibiting hyaluronan production with specific medicine like hymecromone can provide new strategies for effective COVID-19 treatment.

## Acknowledgments

Modified Cas13d plasmid was a generous gift from Professor Pengyu Huang at ShanghaiTech University. We thank Yue Yu for editorial help and comments on the manuscript. This research was supported by Major Special Projects of Basic Research of Shanghai Science and Technology Commission (Grant No. 18JC1411101) the National Natural Science Foundation of China (Grant No. 31872814), and the National Key R&D Program of China (Grant No. 2018YFC1005004).

## Author Contributions

Conceptualization, W.Y.; Formal Analysis, W.L., B.J., S.Y., P.X., Y.T.; Investigation, S.Y., Y.T., L.C., D.Z., P.X., D.R., B.Z., M.L., C.L., C.C., X.R., Y.L., Z.Y., W.L., S.W.; Resources, J.X., H.Z., Z.X., A.L., W.F., S.Y.; Data Curation, W.L., S.Y., P.X., Y.T.; Writing – Original Draft, P.X., S.Y.; Writing – Review & Editing, P.X., S.Y., W.Y., H.W., J.X.; Supervision, W.Y.; Project Administration, W.Y., H.W., J.X.; Funding, W.Y.

## Supplemental Figure Titles and Legends

**Figure S1.**
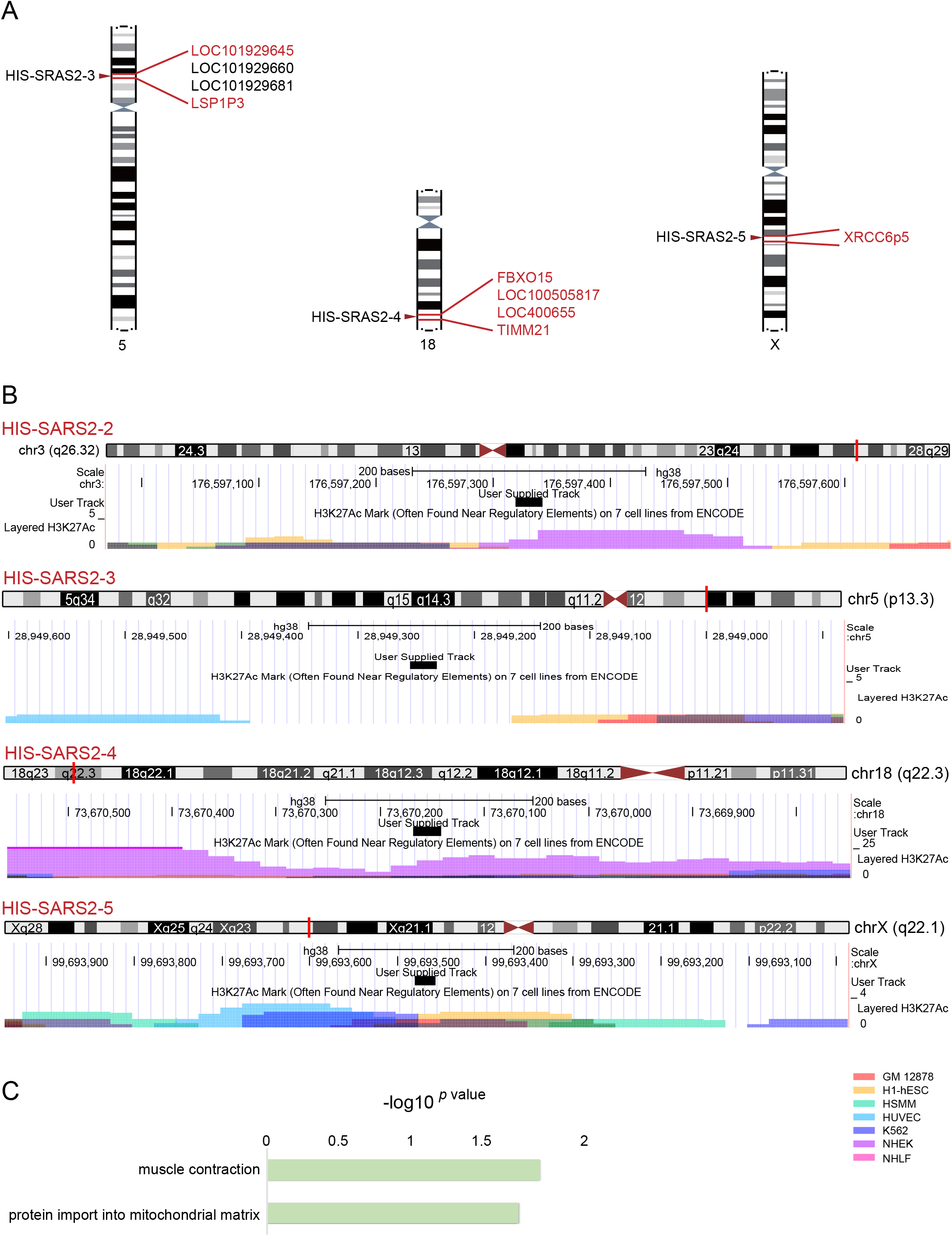
Identification of HIS in SARS-CoV-2. **(A)** The location of the identical sequence of HIS-SARS2-3, HIS-SARS2-4, and HIS-SARS2-5 in human genome and their surrounding genes. All genes within ± 500 kb of those loci are listed. Among them, inflammation- or immunity-related genes, which are defined by text mining (search combining keywords “gene name + inflammation” or “gene name + immunity”) in PubMed and Google Scholar, are written in red. The searched results were further confirmed by literary research. **(B)** The distribution of enhancer marker H3K27ac across the identical sequence of HIS-SARS2-2, HIS-SARS2-3, HIS-SARS2-4, and HIS-SARS2-5 in human genome in seven human cell lines. The upper panel illustrates the corresponding chromatin, and the lower panel illustrates H3K27ac enrichment, within which the location of identical sequence of HIS-SARS2-2/3/4/5 was marked in red line and black block, respectively. **(C)** Gene Ontology (GO) functional annotation analysis of genes within ± 500 kb of the five segments conserved in SARS-CoV-2 and human genomes. The *x*-axis indicates the degree of GO annotation by rich factor. The *y*-axis indicates the functional terms.

**Figure S2.**
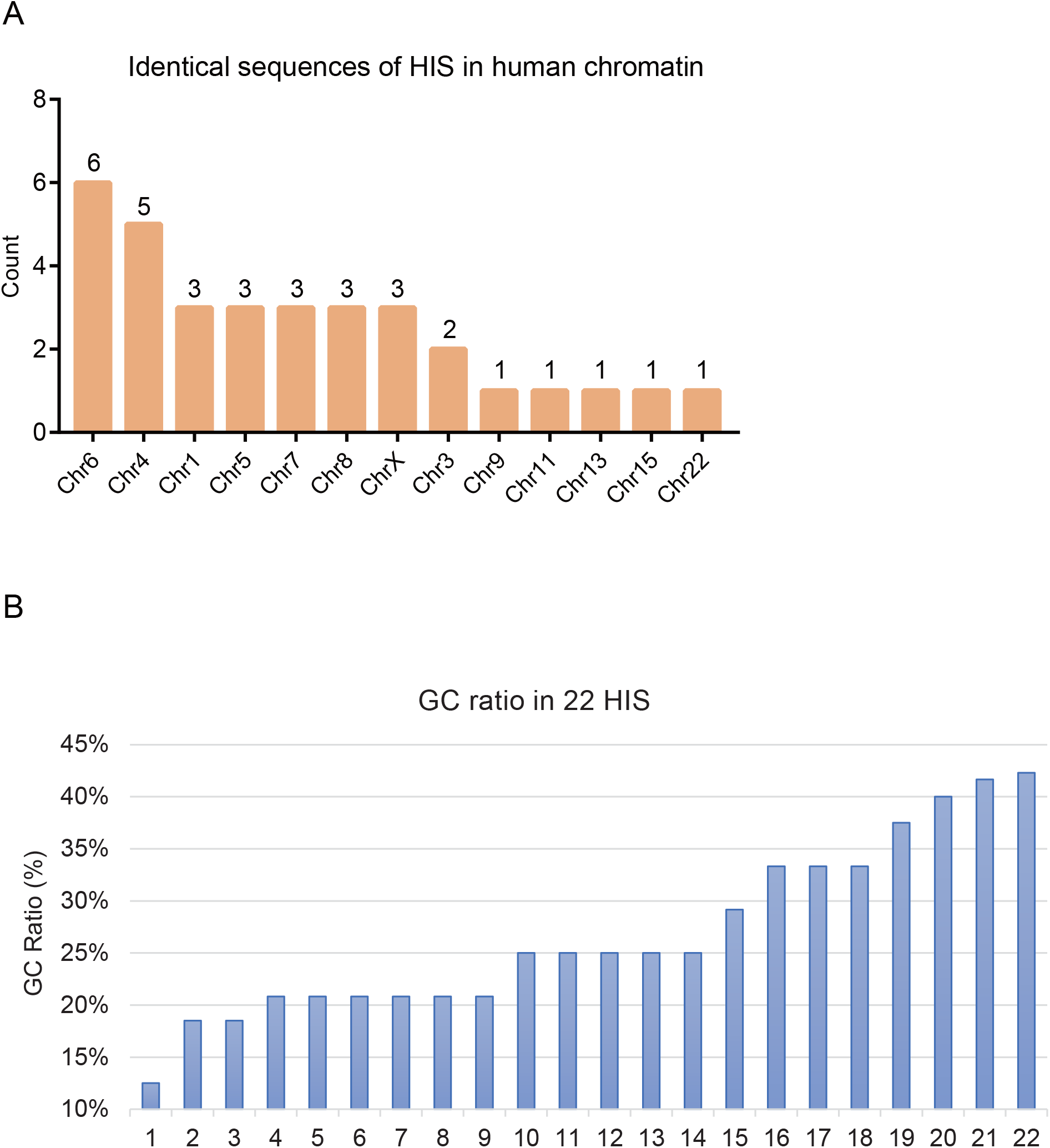
Features of HIS-SARS-CoV-2. **(A)** HIS enrichment in chromatin. **(B)** GC ratio of HIS. The *x*-axis indicates 22 HIS. The *y*-axis indicates the GC ratio (%).

**Figure S3.**
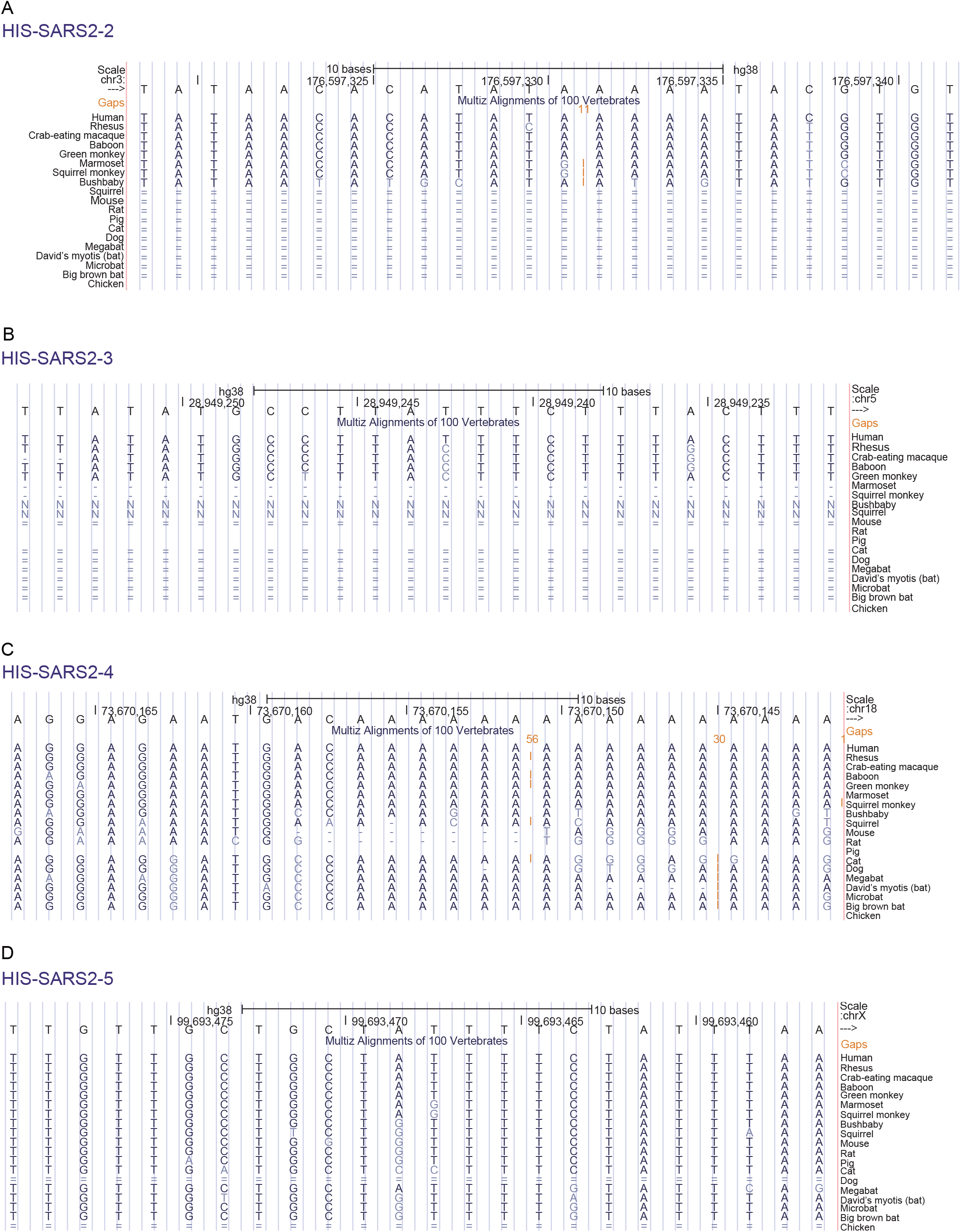
The Conservation of HIS among Species. **(A)(B)(C)(D)** The sequence conservation of HIS-SARS2-2 (A), HIS-SARS2-3 (B), HIS-SARS2-4 (C), and HIS-SARS2-5 (D) in 19 species. The color of bases represents the conservation degree, dark blue represents the fully conserved, while light blue represents the mismatched base, single line (-) indicates no bases in the aligned species, and double-line (=) indicates aligning species has one or more unalignable bases. Ns reflect uncertainty in the relationship between the DNA of both species, due to lack of sequence in relevant portions of the aligning species.

**Figure S4.**
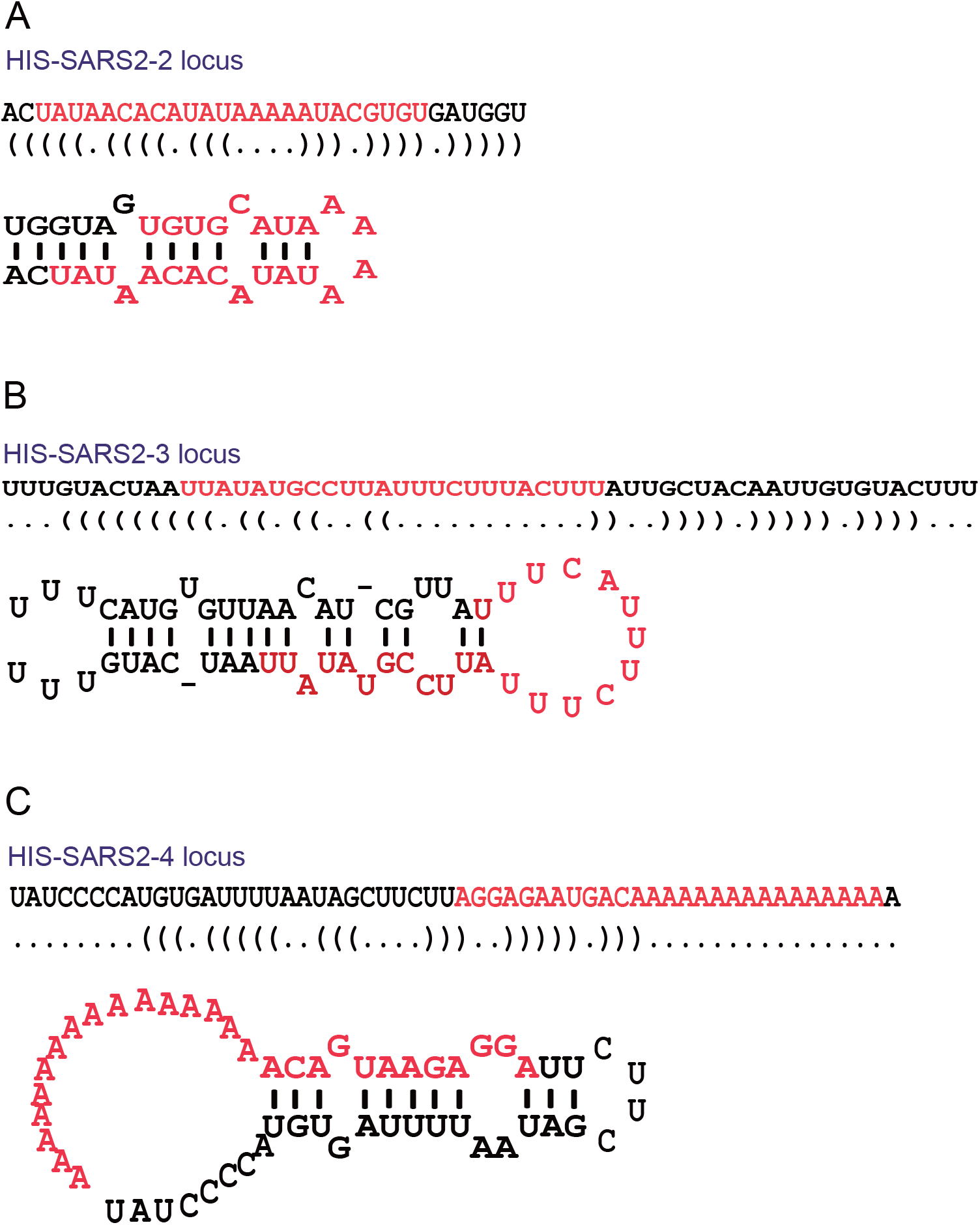
HIS are Predicted to Form miRNA Precursor. **(A)(B)(C)** The predicted RNA secondary structures of HIS-SARS2-2 (A), HIS-SARS2-3 (B), HIS-SARS2-4 (C) and their adjunct bases by the algorithm of minimum free energy. The sequences in red represent HIS-SARS2-2/3/4 RNA.

**Figure S5.**
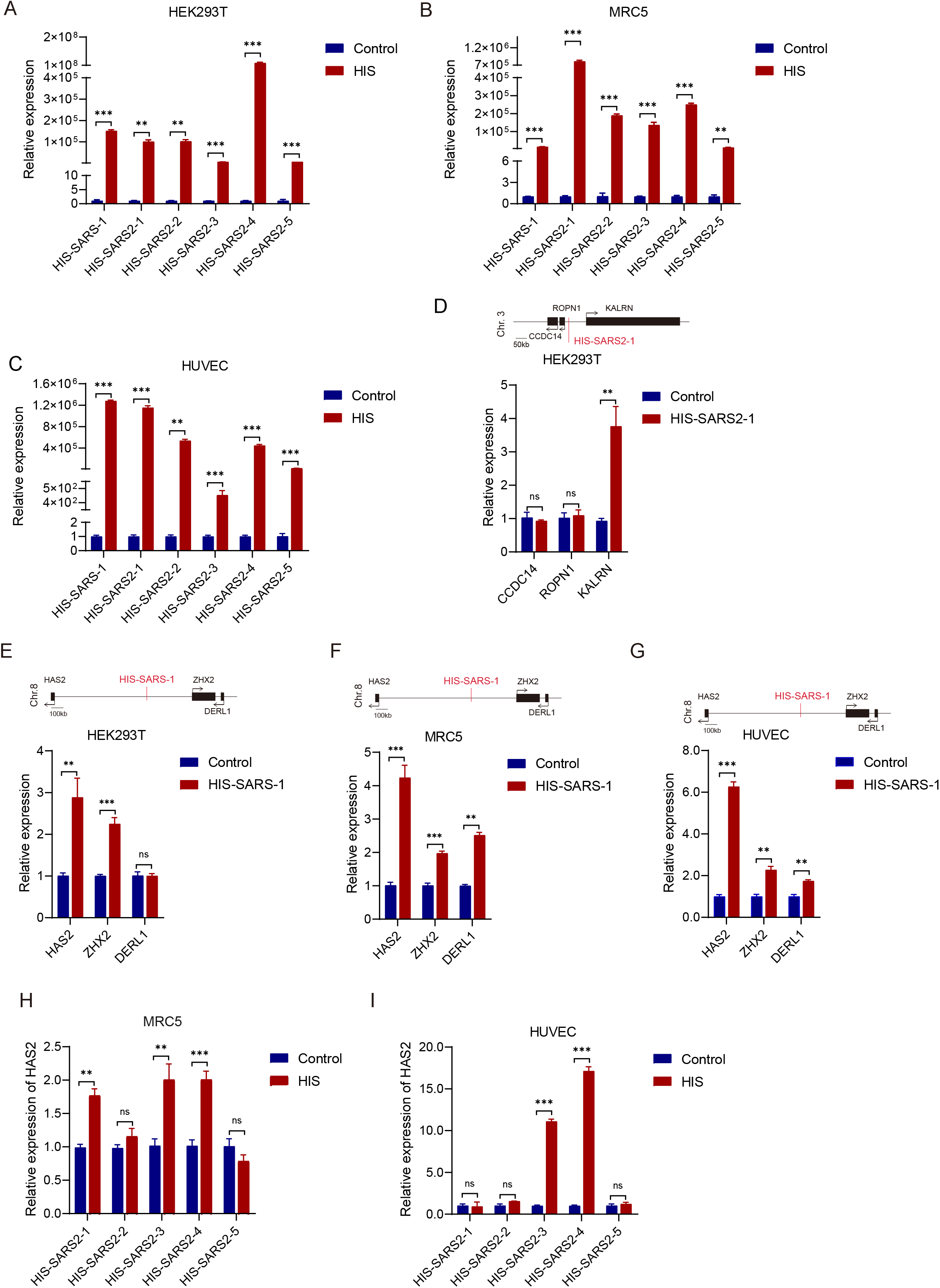
HIS of SARS-CoV and SARS-CoV-2 Upregulated Genes Related to COVID-19 Pathogenic Processes. **(A)(B)(C)** The relative mRNA expression of HIS-SARS-1, HIS-SARS2-1, HIS-SARS2-2, HIS-SARS2-3, HIS-SARS2-4, HIS-SARS2-5 in HEK293T cells (A), MRC5 cells (B) or HUVEC cells (C) after transfected with the corresponding overexpression vectors. **(D)** The relative mRNA expression of the neighboring gene *KALRN* after HIS-SARS2-1 vector transfected in HEK293T cells. **(E)(F)(G)** The relative mRNA expression of the neighboring genes *HAS2*, *ZHX2*, and *DERL1* after HIS-SARS-1 vector transfected in HEK293T cells (E), MRC5 cells (F) or HUVEC cells (G). **(H)(I)** The relative mRNA expression of hyaluronan synthase *HAS2* after five HIS-SARS2 vectors transfected in MRC5 cells (H) or HUVEC cells (I). The *y*-axis indicates the RNA level detected by RT-qPCR. *P*-values were calculated using the unpaired, two-tailed Student’s t test by GraphPad Prism 7.0. *, *P*<0.05; **, *P*<0.01; ***, *P*<0.001; ns, not significant.

**Figure S6.**
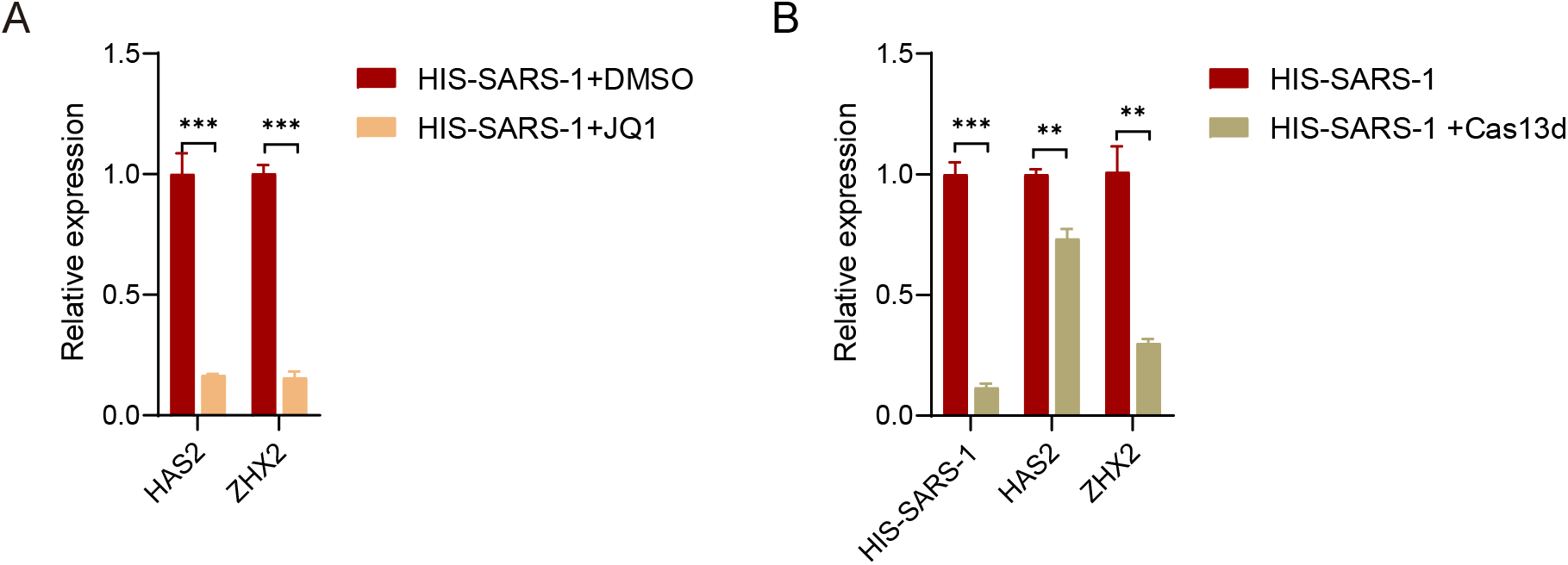
Interference of H3K27ac and Knockdown HIS Abolished the Upregulation of HIS Targeted Genes. **(A)** JQ1 abolished the upregulation of HIS-SARS-1 targeted genes *HAS2* and *ZHX2*. **(B)** Knockdown HIS-SARS-1 by Cas13d abolished the upregulation of HIS-SARS-1 targeted gene *HAS2* and *ZHX2*. Note: The *y*-axis indicates the RNA level detected by RT-qPCR. *P*-values were calculated using the unpaired, two-tailed Student’s t test by GraphPad Prism 7.0. **, *P*<0.01; ***, *P*<0.001.

**Figure S7.**
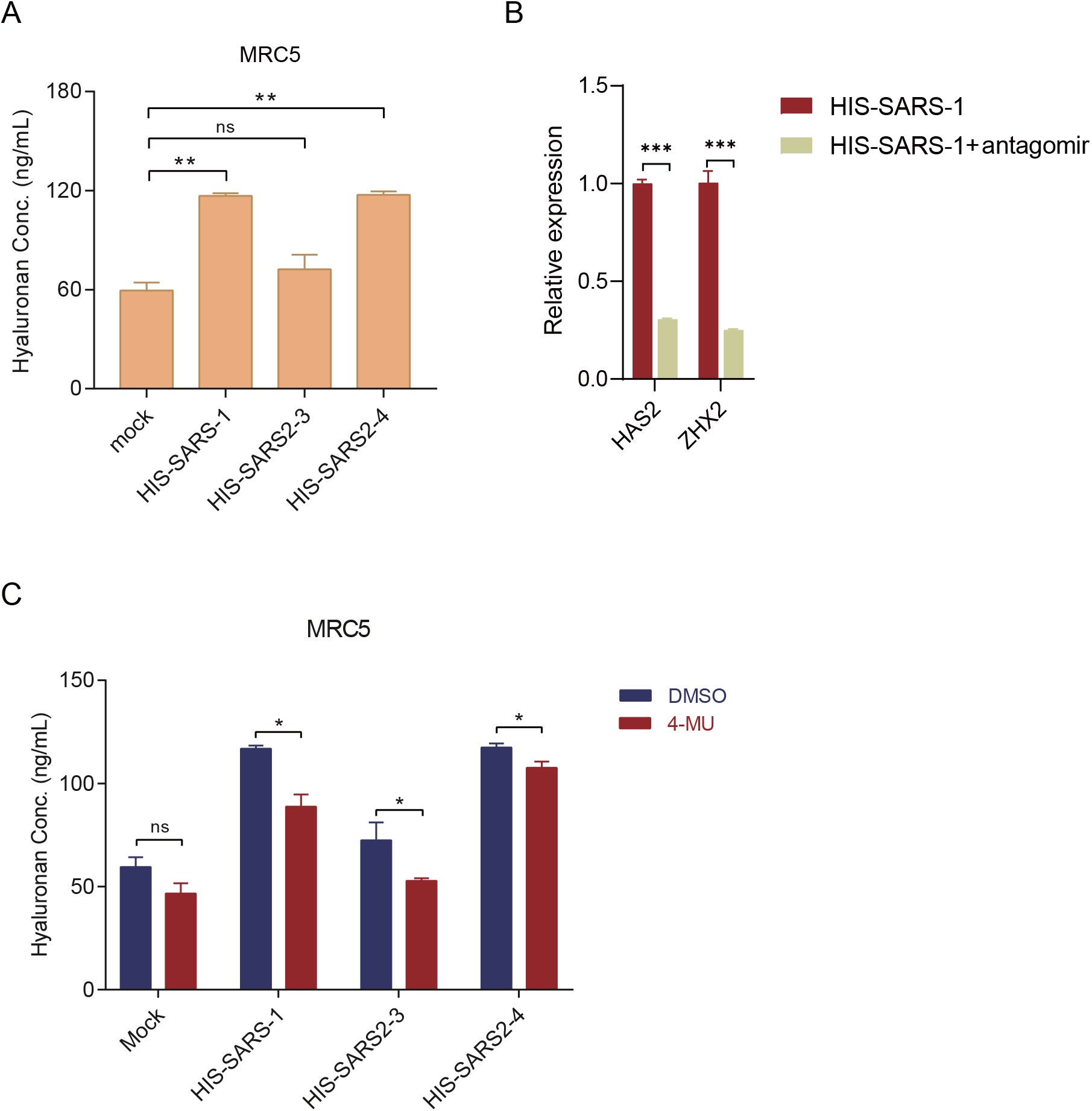
Therapeutic Potentials of Targeting Hyaluronan. **(A)** Hyaluronan released in cell culture supernatants of Mock, HIS-SARS-1, HIS-SARS2-3, and HIS-SARS2-4 overexpressed in MRC5. The *y*-axis indicates the hyaluronan level detected by ELISA. *P*-values were calculated using the unpaired, two-tailed Student’s t test. **, *P*<0.01. **(B)** Antagomir of HIS-SARS-1 blocks the upregulation of HIS-SARS-1 targeted gene *HAS2* and *ZHX2*. The *y*-axis indicates the RNA level detected by RT-qPCR. *P*-values were calculated using the unpaired, two-tailed Student’s t test by GraphPad Prism 7.0. ***, *P*<0.001. **(C)** ELISA for HA release on cell culture supernatants of mock, HIS-SARS-1, HIS-SARS2-3, and HIS-SARS2-4 stably expressed in MRC5 treated with 100 μM 4-MU and DMSO as control. The *y*-axis indicates the hyaluronan level detected by ELISA. *P*-values were calculated using the unpaired, two-tailed Student’s t test. ns, not significant; *, *P*<0.05.

## Supplemental Tables

Supplemental Table 1. The accession numbers

Supplemental Table 2. HIS in HCoVs

Supplemental Table 3. Sequences used in the study

Supplemental Table 4. Human identical sequences in multiple viruses

Supplemental Table 5. Overlapped genes regulated by HIS-SARS2 in lung

## STAR★METHODS

**KEY RESOURCES TABLE**

| REAGENT or RESOURCE | SOURCE | IDENTIFIER |
| --- | --- | --- |
| Antibodies |  |  |
| H3K27ac antibody (ChIP Grade) | abcam | cat# ab4729 |
| Rabbit IgG (ChIP Grade) | abcam | cat# ab171870 |
| Bacterial and Virus Strains |  |  |
| DH5α | TIANGEN biotech | cat# CB101-02 |
| Biological Samples |  |  |
| Plasma of COVID-19 patients | Shanghai Public Health Clinical Center | N/A |
| Chemicals, Peptides, and Recombinant Proteins |  |  |
| Puromycin | Sangon Biotech | A610593-0025 |
| VAHTS DNA Clean Beads | Vazyme | cat# N411-01 |
| BioMagPlus Concanavalin A | Bangs Laboratories | cat#BP531 |
| JQ1 | Selleck Chemicals Inc. | S7110 |
| 4-Methylumbelliferone (4-MU) | Selleck Chemicals Inc. | S2256 |
| Critical Commercial Assays |  |  |
| ClonExpress II One Step Cloning kit | Vazyme | cat#C112 |
| Hyaluronan DuoSet ELISA | R&D Systems | DY3614-05 |
| DuoSet ELISA Ancillary Reagent Kit 3 | R&D Systems | DY009 |
| Deposited Data |  |  |
| Data of this study | Mendeley Data | https://data.mendeley.com/datasets/853mrdr9cs/draft?preview=1 |
| Experimental Models: Cell Lines |  |  |
| HEK293T | Our lab | N/A |
| MRC5 | Our lab | N/A |
| HUVEC | ALLCELLS | HUVEC-001F |
| Oligonucleotides |  |  |
| Antagomirs for HIS, see Table S3 | This paper | N/A |
| Primers for HIS and target genes, see Table S3 | This paper | N/A |
| sgRNAs targeting pre-HIS, see Table S3 | This paper | N/A |
| Sequences in EMSA, see Table S3 | This paper | N/A |
| Recombinant DNA |  |  |
| pSPAX2 | Our lab | N/A |
| pMD2G | Our lab | N/A |
| pCDH-CMV-MCS-EF1-copGFP | This paper | N/A |
| Modified Cas13d plasmid | Prof. Pengyu Huang at ShanghaiTech Univ. | N/A |
| Software and Algorithms |  |  |
| GraphPad Prism (version 7.00) | GraphPad Software | https://www.graphpad.com/ |
| Adobe Illustrator CS6 (version 16.0.0) | Adobe | https://www.adobe.com/ |
| MiRanda (3.3a) | Betel et al., 2010 | http://www.miranda.org/ |
| Other |  |  |
| Hieff Trans<sup>TM</sup> liposome transfection reagent | YEASEN | 40802ES02 |
| SYBR Green | TIANGEN | FP205-02 |
| TRIzol Reagent | Invitrogen | 15596018 |
| PrimeScript<sup>TM</sup> RT reagent Kit with gDNA Eraser | TAKARA | RR047A |

### Lead Contact

Further information and requests for resources and reagents should be directed to and will be fulfilled by the Lead Contact, Wenqiang Yu.

### Materials Availability

This study did not generate new unique reagents.

### Data and Code Availability

All the data are provided as Figures, supplementary Figures, or supplementary tables. Data alongside this manuscript have been deposited in Mendeley Data and the preview linkage, https://data.mendeley.com/datasets/853mrdr9cs/draft?preview=1. The parameters used in the analysis refers to Method Details.

## EXPERIMENTAL MODEL AND SUBJECT DETAILS

### Cells cultures

HEK293T (human embryonic kidney transformed cell) and MRC5 (human fetal lung fibroblast cells) cells were cultured in DMEM/High glucose medium (HyClone) supplemented with 10% FBS (Gibco) and 1% Penicillin-Streptomycin solution (HyClone). HUVEC (human umbilical vein endothelial cell, bought from AllCells) was cultured in the commercial culture medium (AllCells, H-004) according to the manufacturer’s guidelines.

### Clinical subjects

We retrospectively analyzed 137 COVID-19 patients admitted to The Shanghai Public Health Clinical Center (SPHCC), which was approved by the SPHCC Ethics Committee. These cases were confirmed in strict accordance with the World Health Organization diagnostic criteria. All participants were provided with written informed consent for sample collection and subsequent analyses. According to the Diagnosis and Treatment Protocol for Novel Coronavirus Pneumonia (Trial Version 7) released by National Health Commission & State Administration of Traditional Chinese Medicine on March 3, 2020, COVID-19 patients with pneumonia were categorized as mild and severe based on the characteristic pneumonia features of chest CT. During the hospitalization of COVID-19 patients, they accepted necessary laboratory examinations and imaging examinations at any time according to their conditions, including lymphocyte count (137 patients), D-dimmer (132 patients), and C-reactive protein (CRP, 135 patients). Besides, ELISA was performed to detect the hyaluronan level in their plasma at the same time points.

## METHOD DETAILS

### Viral Genome and Human Genome Blast

Human reference genome (hg38) was obtained from GenBank using Blastn (Chen et al., 2015) with SARS-CoV-2 as a query. The detail parameters were chosen as follow: 1) The Maximum number of hits to report:500; 2) Maximum E-value for reported alignments: 10; 3) Word size for seeding alignments: 11; 4) the match/Mismatch scores: 1,-3; 5) Gap penalties: Openging:2, Extension:2; 6) Filter low complexity regions; 7) Filter query sequences using RepeatMasker.

### Gene Function Annotation

KEGG (Kyoto Encyclopedia of Genes and Genomes) and Gene Ontology analysis on the surrounding (±500kb) genes of the identical sequences of HIS in human genome were performed using DAVID (Huang da et al., 2009). Gene ontology Biological Processes and KEGG pathway were ranked by *P*-value and the top 10 terms were plotted.

### Plasmid Construction and Antagomir Synthesis

Five HIS in SARS-CoV-2 were chosen to construct plasmids, named HIS-SARS2-1, HIS-SARS2-2, HIS-SARS2-3, HIS-SARS2-4, and HIS-SARS2-5. Besides, we chose HIS-SARS-3 in SARS-CoV as the parallel group. In brief, as HIS precursors, 100~150bp virus DNA fragments containing 24~27 bp HIS in SARS-CoV and SARS-CoV-2 were obtained by annealing and extension with specific primers synthesized by Shanghai SunnyBio-technology Co., Ltd. Then, these HIS were cloned into the pCDH-CMV-MCS-EF1-copGFP lentiviral vector (Zou et al., 2018) at EcoR I(5′) and BamH I (3′) sites through ClonExpress II One Step Cloning Kit (Vazyme, C112) according to the manufacturer’s manual. Besides, diverse single guide RNAs (sgRNAs) targeting the HIS precursors were designed and cloned into modified lentivirus plasmids containing Cas13d. Also, the antagomirs for HIS-SARS2-1, HIS-SARS2-2, HIS-SARS2-3, HIS-SARS2-4, HIS-SARS2-5, and HIS-SARS-1 were purchased from Guangzhou RiboBio Co., Ltd. The sequences of primers, HIS precursors, and antagomirs are listed in TableS3.

### Vector Transfection

Cells were transfected with plasmids or antagomirs at 70%-80% confluency via Hieff Trans^TM^ liposome nucleic acid transfection reagent (YEASEN, 40802ES02) following the manufacturer’s instruction. We changed fresh medium containing 10% FBS at 6-8 hours after transfection and harvested transfected cells at 72 hours after transfection to detect the expression of HIS and target genes or perform ChIP assay.

### Lentivirus Package and Cell Screening

We co-transfected pCDH-pre-HIS, pSPAX2, and pMD2G plasmids into HEK293T cells in a ratio of 4:3:1.2 and collected the virus supernatant by filtering cell culture supernatant with 0.45 filters at 48 hours after changing the serum-containing medium. Then, cells were infected with different lentiviruses and cultured in medium with 1μg/ml puromycin to obtain stable cell lines.

### Quantitative RT-PCR (RT-qPCR)

Total RNA was extracted from freshly harvested cells using TRIzol Reagent (Invitrogen, 10296028). Complementary DNA (cDNA) was synthesized with the PrimeScript™ RT reagent Kit (Takara, RR047A) involved in genomic DNA erasing. Quantitative PCR was performed using SYBR Green Pre-Mix (TIANGEN, FP205) on the Roche LightCycler480 instrument. GAPDH was the normalized endogenous control gene. Relative gene expression was calculated according to 2^−ΔΔCt^ method. The primers for target genes and diverse HIS precursor fragments are shown in TableS3.

### Chromatin Immunoprecipitation (ChIP)

ChIP assay was carried out as our previous study described(Xiao et al., 2017). In brief, transfected cells were cultured in 10cm dishes and crosslinked with 1% formaldehyde in 1×PBS for 10 minutes at room temperature. After sonication, sheared chromatin was immunoprecipitated with H3K27ac antibody (Abcam, ab177178) and Protein A magnetic beads (Invitrogen, 10002D) overnight at 4°C. DNA from the chromatin immunocomplexes was extracted with QIAquick PCR Purification Kit (QIAGEN, 28106) according to the manufacturer’s guidelines. ChIP-derived DNA was analyzed by quantitative PCR using SYBR Green PreMix (see TableS3 for primer sequences), and data were normalized by input DNA.

### Loss-of-function Assay for HIS

Loss-of-function of HIS was performed by inhibiting the enhancer components with JQ1 (Selleck Chemicals Inc., S7110) as previously described (Suzuki et al., 2017) or blocking them via antagomirs synthesized by Guangzhou RiboBio Co., Ltd (see TableS3 for antagomir sequences). Cells were co-transfected with antagomirs and HIS precursor vectors and harvested at 72 h. Besides, cells transfected with HIS precursor vectors were then treated with 500 nM JQ1 for 24 h. As indicated time points, total RNA was exacted from the harvested cells and evaluated the effect of loss-of-function for HIS on the target genes expression by RT-qPCR.

### UV-Vis Spectroscopy Measurement

UV-Vis spectroscopy experiments were performed on a SHIMADZU UV-1900 UV-Vis spectrophotometer (Tokyo, Japan) at room temperature. The DNA or/and RNA oligonucleotide, DNA-RNA hybrid, DNA duplex samples (2 mL) were diluted to 2.0 μM in 10 mM Tris-HCl, 50 mM NaCl buffer at pH 7.0. Spectra were recorded over a wavelength range from 350 to 200 nm with a scan rate at 100 nm/min and data interval for 1 nm. A 10 mm optical path length quartz cuvette was used for UV-Vis measurement.

### Thermal Melting (T_m_) Analysis

Melting temperatures (Tm) of self-complementary sequences were determined from the changes in absorbance at 260 nm as a function of temperature in a 10 mm path length quartz cuvette on a SHIMADZU UV-1900 UV-Vis spectrophotometer (Tokyo, Japan) equipped with a temperature control system. Solutions (2 mL) of 2 μM pre-hybridized DNA-RNA hybrid or DNA duplex in aqueous buffer (10 mM Tris-HCl, 50 mM NaCl, pH 7.5) were equilibrated at 10°C for 5 min and then slowly ramped to 90°C with 2°C step at a rate of 1°C/min. Tm values were calculated as the first derivatives of heating curves.

### Probes Preparation

All DNA-RNA hybrid and DNA duplex were prepared as follows: Firstly, the oligonucleotide was mixed with an equal molar of the complementary target strand in hybridization buffer (10 mM Tris-HCl pH 7.8, 50 mM NaCl, 1 mM EDTA), then the mixtures were annealed by heating them to 95°C for 5 min, and then slowly cooled to room temperature. Secondly, the annealed mixtures were separated on 16% nondenaturing polyacrylamide gel at 100 V for 120 min using 0.5×TBE as electrophoresis buffer. Thirdly, the target bands in the gel were cut and recovered the nucleic acid probes from the gel according to the reference [Molecular Cloning: A Laboratory Manual, Fourth Edition, by Green MR, Sambrook J, 2012 Cold Spring Harbor Laboratory Press, Cold Spring Harbor, New York, USA].

### Protein Purification

Full-length human argonaute 2 (hAGO2) coding sequence was amplified and cloned into the BamH I and Hind III restriction endonuclease sites of a home-reconstructed pMAL-C5X expression vector for protein purification. The constructed plasmid was transformed into a laboratory-built Dam knock out Rosetta (DE3) cells for protein expression. The colony was inoculated into 50 mL of LB containing 100 μg/mL ampicillin and allowed to grow overnight at 37°C. This culture was diluted into 4L LB containing 100 μg/mL ampicillin and grown at 37°C until A600 reached 0.4~0.6. Then the culture was overnight induced by the addition of 0.2 mM isopropyl-1-thio-β-D-galactopyranoside at 16°C. The cells were harvested at 10 000 rpm for 5 min at 4°C and then suspended in 100 mL lysis buffer (20 mM Tris-HCl pH 7.5, 500 mM NaCl, 10% glycerol, 1 mM DTT) supplemented with EDTA-free protease inhibitor. The suspended cells were lysed by ultrahigh-pressure continuous flow cell breaker under the low-temperature (4°C) water bath. Following centrifugation 12,000 rpm for 30 min at 4°C, the cleared lysate was loaded onto 5 mL MBP TrapTM HP column pre-equilibrated with 20 mM Tris-HCl pH 7.5, 200 mM NaCl, 10% glycerol, 0.2 mM DTT. The MBP-hAGO2 recombinant protein was eluted by the 20 mM Tris-HCl pH 7.5, 200 mM NaCl, 1.0 mM maltose, 10% glycerol, 0.2 mM DTT. The eluted recombinant protein was digested by thrombin to remove the MBP-tag under the ice-water bath, and then the digested mixture was further purified by the 1 mL HisTrap HP column.

### Electrophoretic Mobility Shift Assay (EMSA)

The oligonucleotide probes (20 nM) were incubated with hAGO2 recombinant protein on the ice for 30 min in the fresh prepared binding buffer (10 mM PBS pH 7.5, 5 mM MgCl_2_, and 0.1% Triton-100). The protein-substrate complexes were separated from the unbound substrate probes on 5% nondenaturing polyacrylamide gels at 100 V for 50min using 0.5×TBE as electrophoresis buffer. After electrophoresis, the resolved oligonucleotide probes in the gel were detected using an Odyssey CLx dual-color IR-excited fluorescence imaging system (LI-COR, Lincoln, NE).

### Enzyme-linked Immunol Sorbent Assay (ELISA)

Hyaluronic acid of COVID-19 patients’ plasma was measured in 1:5 dilution by the enzyme-linked sandwich assay Hyaluronan DuoSet ELISA (R&D Systems, Minneapolis, MN, USA) following the manufacturer’s descriptions. To evaluate the effect of hyaluronic acid inhibitor, 5 × 10^6^ cells were seeded in 12-well plates and incubated for 24 h under appropriate treatments (with 100 μM 4-MU or 200 μg/ml hymecromone, and DMSO as the control group). Then, we collected the culture supernatants and quantified hyaluronic acid using the same ELISA kit.

## QUANTIFICATION AND STATISTICAL ANALYSIS

The mRNA relative expression level for each sample was evaluated by 2^−ΔΔCt^ method. The statistical analysis was described in figure legends.

